# Altered SNr activity under dopamine depletion: mechanistic predictions from data-driven network models

**DOI:** 10.1101/2025.08.15.670540

**Authors:** John E. Parker, Asier Aristieta, Ya Emma Gao, Aryn H. Gittis, Jonathan E. Rubin

## Abstract

Sufficient loss of dopamine within the basal ganglia (BG) leads to neuronal activity changes, including altered firing rates and firing patterns, thought to underlie parkinsonian motor symptoms. Yet, within BG neuronal populations, baseline activity and responses to inputs are highly variable, complicating efforts to identify key factors associated with pathological changes. We introduce a novel approach to constructing a computational neuron population model that, when applied to the mouse substantia nigra pars reticulata (SNr), captures the firing heterogeneity observed across slice and *in vivo* recordings. This model reproduces the diversity of SNr neuron responses to stimulation of GABAergic input terminals, yielding new insights into the mechanisms underlying this variability. Moreover, our modeling pinpoints significant decreases in TRPC3 conductance in SNr dendrites as a key determinant of altered SNr activity in the dopamine depleted state, with important implications for efforts to restore functional SNr activity in this condition.

**Teaser:** In silico model capturing heterogeneity suggests factors behind SNr altered activity in dopamine depletion and responses to input

## Introduction

The basal ganglia (BG) are considered as a subcortical hub for reinforcement learning and action selection (*1–3*). The classical theoretical framework casts the outputs from the BG to the rest of the brain, which are inhibitory, as a gate, with suppression of a subset of these outputs needed to “open the gate” and allow a selected action to proceed (*4, 5*). In conditions of depletion of the neurotransmitter dopamine, such as in Parkinson’s disease, the gate is believed to be strengthened, resulting in suppressed activation of motor pathways. A variety of experimental findings, however, point to a more complicated situation. One example of this complexity arises from stimulation of striatal neurons that project, either directly or through multiple synapses, to BG output nuclei. Targeted stimulation of striatal spiny projection neurons (SPNs) that express the D1 dopamine receptor can have strongly pro-kinetic effects (*6*); however, the responses of individual BG output neurons to this stimulation is diverse, with some neurons showing the reduction in activity predicted by the classical model but others showing no effect or even increased activity (*7, 8*). Similarly, targeted stimulation of D2-receptor expressing SPNs that leads to anti-kinetic effects also yields diverse BG output neuron responses, as does stimulation of globus pallidus neurons that are classically expected to inhibit BG outputs (*9, 10*).

Since many therapeutic interventions for Parkinson’s disease are based on the goal of mitigating pathological BG outputs, and long-term motor rescue in dopamine depleted mice is associated with changes in BG output neuron activity (*11*), it is critically important to elucidate the factors that affect these outputs and how they change under dopamine depletion. This represents a challenging aim, however, since BG output neuron activity and responses to inputs are so heterogeneous and emerge in a network of neurons that receive inputs from multiple sources and that interact directly through recurrent collaterals. To address this challenge, we have previously turned to computational modeling (*8, 12*). By constructing biologically principled models of individual and interacting neurons in the substantia nigra pars reticulata (SNr), the major motor output of the rodent BG, we identified the potential of variations in the ongoing inhibitory tone and the associated intracellular chloride concentration of SNr neurons to account for their diverse responses to similar inputs and to induce the emergence of oscillations in the delta frequency band, a key signature of motor complications in dopamine depleted rodents (*13*).

Despite these insights, the relative contributions of chloride-related effects versus other factors to the input responses of SNr neurons remain unknown. Moreover, separate experiments have suggested a range of possible changes in SNr neuron inputs and intrinsic properties that may arise with dopamine depletion, and how these effects translate into altered SNr responses, and hence BG outputs, has not been explored. These issues led us to ask whether we could achieve further advances by incorporating a rich set of experimental recordings of SNr activity into a computational model for a network of synaptically interconnected SNr neurons. Standard neuronal modeling approaches involve using experimental data to identify the key trans-membrane ion currents present in the neurons of interest and to estimate parameters associated with these currents. A network model can be constrained by additionally measuring synaptic properties and diversifying the population of neurons in the model network by choosing parameter values from distributions centered around the original estimates. These steps, unfortunately, are limited by a lack of data about the nature of parameter distributions and correlations among parameters in real neurons, especially considering findings that quantities such as ionic conductance can vary widely across neurons within a single population that produce similar outputs (*14, 15*).

In this work, we present results obtained using a new computational modeling approach in which we harness the full complexity of a large collection of neuronal unit recordings, obtained across multiple conditions, to build a model network customized to match the data (Figure 1). Specifically, we generate a large database of potential model neurons and select units from this database with firing properties that match individual unit recordings. We subsequently assemble our selected model neurons into a model network, with experimentally measured coupling properties, and consider how the distributions of the coupled neurons’ firing properties match what we observed in units in the corresponding biological network. While performing this procedure based on a single data set would risk overfitting, we validate our model network’s performance across not just baseline conditions but also two forms of optogenetic stimulation. As a result of this customized network construction, we were able to achieve two major advances. First, we could make predictions about the relative contributions of disinhibition, chloride effects, and other factors in shaping the distribution of SNr responses to stimulation of the SNr’s primary GABAergic input pathways. Second, and most excitingly, we could also generate predictions about what cellular and network features in the SNr are most likely to underlie altered SNr activity under dopamine depletion. These predictions could be valuable for guiding future work to restore BG function after pathological dopamine depletion occurs.

**Figure 1:**
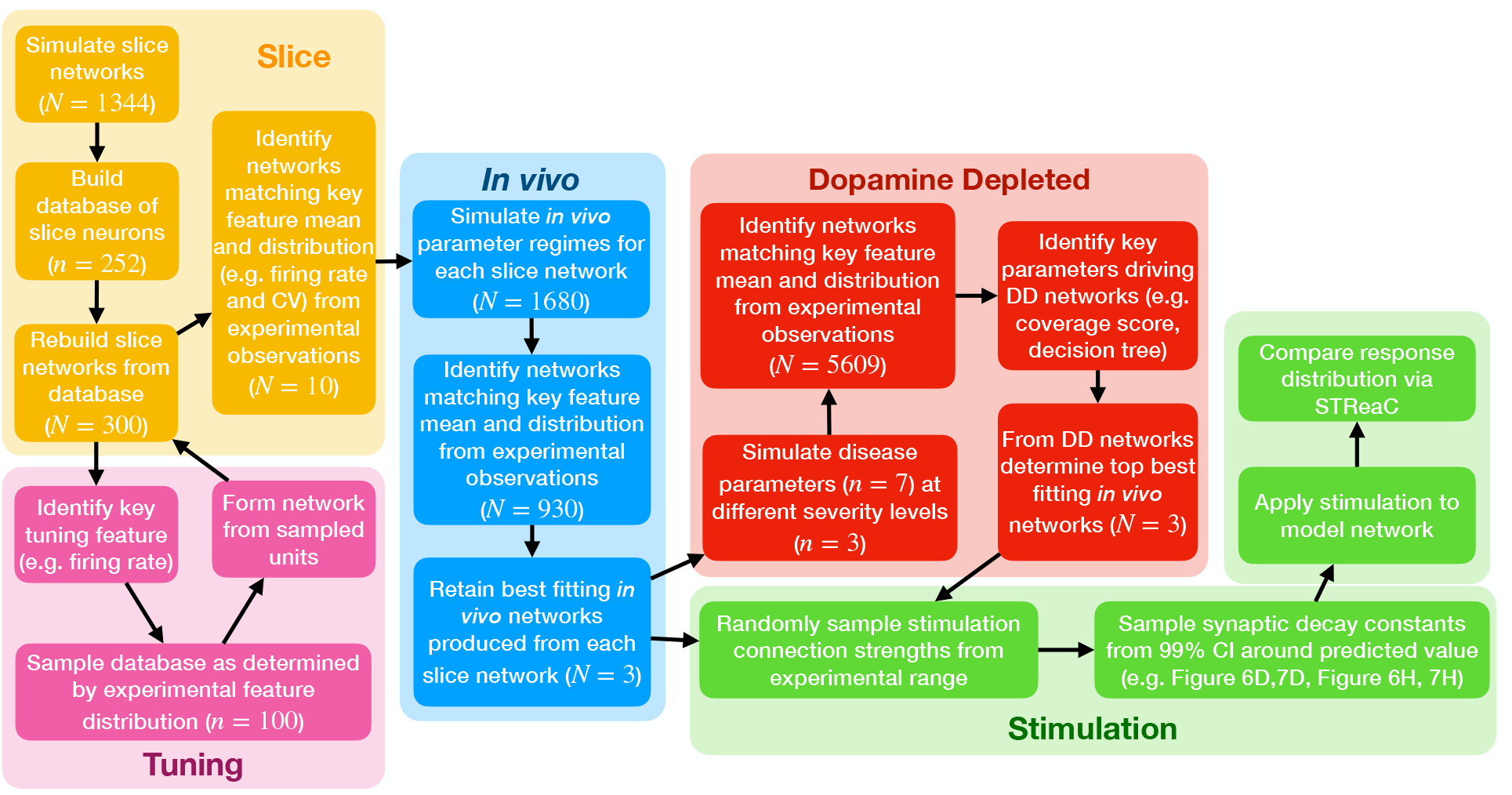
Flowchart of computational modeling pipeline. Our process starts with the steps in the Slice box and proceeds following the arrows. In two cases the arrows are intentionally ambiguous: the third step in Slice involves a detour to the Tuning box before proceeding to the fourth step, and the *In vivo* box can be followed by the stimulation-response analysis described by the Stimulation box or by the DD analysis described in the Dopamine Depleted box, with subsequent analysis of responses to stimulation for the DD networks. In all cases, numbers of networks are designated with *N* and numbers of neurons with *n*.

## Results

### Standard Approach to Tuning SNr Model Network Parameters Fails to Match Slice Recordings

To study the possible factors underlying the diversity of SNr responses to GABAergic inputs and the alterations in SNr firing under dopamine depletion, we constructed an SNr network model in a way that strongly leveraged SNr recordings under a range of conditions (Figure 1). As in an earlier study (*8*), each SNr neuron in the model is a conductance-based unit with two compartments, one to represent the soma where synapses from the external segment of the globus pallidus (GPe) occur and another to represent the dendrite where striatal synapses arise (Figure 2A; see *Methods* for details of equations and a list of parameter values, in Table 1, that have changed relative to earlier work (*8*)), with a baseline parameter set based on experimental constraints that produces a firing rate of approximately 10 Hz in the absence of synaptic inputs.

**Table 1:**
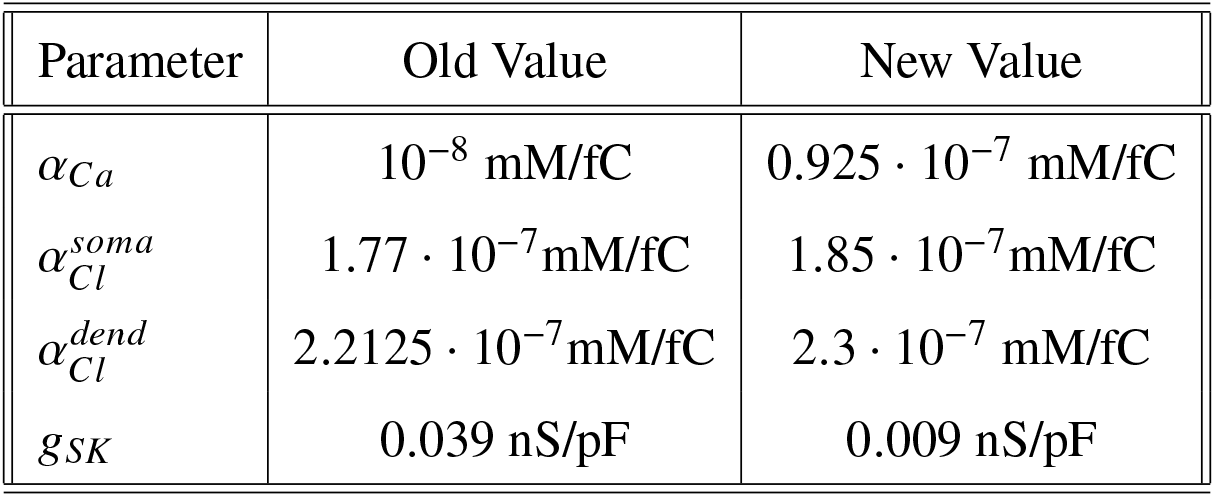
Table of parameter changes from (*8*).

| Parameter | Old Value | New Value |
| --- | --- | --- |
| $\alpha_{Ca}$ | $10^{-8}$ mM/fC | $0.925 \cdot 10^{-7}$ mM/fC |
| $\alpha_{Cl}^{soma}$ | $1.77 \cdot 10^{-7}$ mM/fC | $1.85 \cdot 10^{-7}$ mM/fC |
| $\alpha_{Cl}^{dend}$ | $2.2125 \cdot 10^{-7}$ mM/fC | $2.3 \cdot 10^{-7}$ mM/fC |
| $g_{SK}$ | 0.039 nS/pF | 0.009 nS/pF |

**Figure 2:**
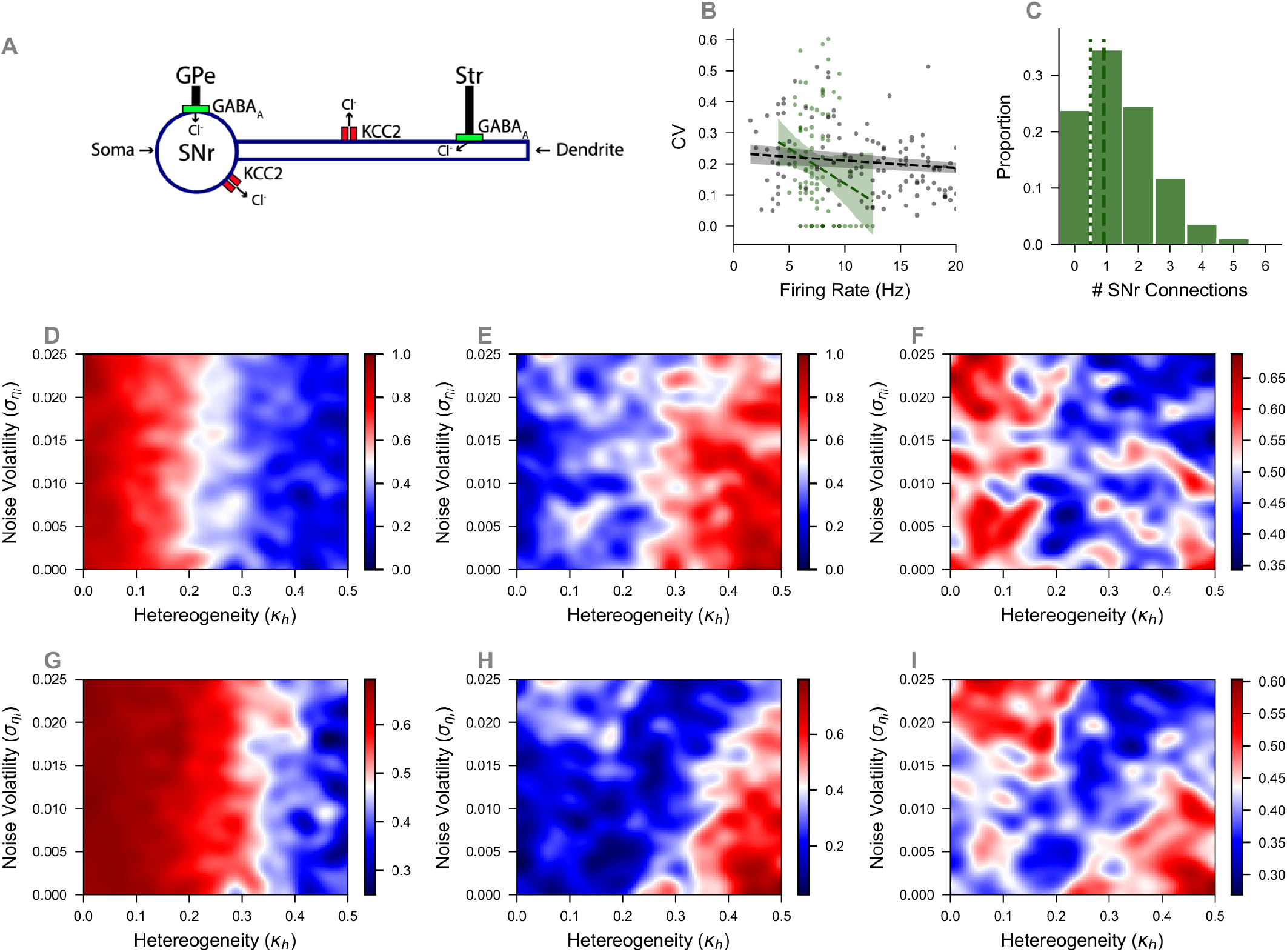
Model SNr neuron and slice network construction. (**A**) Schematic illustration of SNr neuron model from (*8*). (**B**) Linear regressions with 99% confidence intervals for the model (green) and experimental (gray) CV-firing rate relations. Model linear regression has slope = −0.022, intercept = 0.36, and *R*^2^ = 0.047 while experimental linear regression has slope = −0.002, intercept = 0.235, and *R*^2^ = 0.118 (**C**) Connection histogram from across 400 slice networks of 100 neurons each. (**D**) Heatmap of normalized relative error in firing rates between slice model with varying levels of current noise and heterogeneity and experimental data. (**E**) Heatmap of normalized relative error in CV. (**F**) Averages of values in panels D and E. (**G**) Heatmap of relative error between the means of the firing rate distributions from the slice model with varying levels of current noise and heterogeneity and exerpimental data. (**H**) Heatmap of relative error in the mean CV. (**I**) Averages of values in panels G and H.

We initiated the construction of an SNr network model based on recordings in SNr slice preparations, which include fewer complicating factors, and hence free parameters to constrain, than *in vivo* recordings. Thus, we aimed to introduce synaptic coupling in a collection of model SNr neurons to achieve distributions of firing rates and coefficients of variation (CVs) that matched experimental recordings from SNr slices from mice. Specifically, we compared simulated network behavior to two seconds of firing data (the length of our shortest recording) from each of 252 recorded units in the absence of experimental manipulations (e.g., optogenetic stimulation). Although we had recordings from 260 units, we z-scored this data and removed the 8 outlier units for which either firing rate or CV z-score magnitude exceeded 3.

We followed a fairly standard initial approach to constructing our model network. In line with our earlier study (*8*), we worked with a network of 100 model SNr neurons. We introduced synaptic coupling between the model SNr cells with a pairwise connection probability (approximately 0.014) reflecting the reported prevalence of unitary synaptic connections in SNr slice networks (*16*) (average of 1.16 ± 1.07 connections from 81 cells). Connecting 100 identical model SNr neurons in this way led to a distribution of numbers of SNr connections to different neurons but was insufficient to produce the range of firing rates and CVs observed experimentally (Figure 2B). Hence, we next introduced current noise and cell parameter heterogeneity to enhance the biological realism of the model and to attempt to improve its agreement with the experimental data. The current noise represents intrinsic background fluctuations in ion channel opening and in synaptic input currents and was modeled as independent Ornstein-Uhlenbeck (OU) processes (*17–19*) in each of the soma and dendrite equations, with the volatility and mean rate of reversion of the OU process in the dendrite scaled relative to those in the soma to account for capacitance differences. We implemented heterogeneity by sampling each maximal conductance value *x*_*i*_ for each voltage-gated channel type for each neuron from a normal distribution, 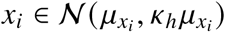, where 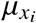 is the baseline value for *x*_*i*_ (*8*) and *k*_*h*_ is a common scaling factor that relates the standard deviation of each distribution to its mean (see Eq. 6 in *Methods*). Additionally, given past evidence of its importance (*8*), we imposed heterogeneity in the GABA reversal potential *E*_*GABA*_ across our model neurons, with positive correlations between neurons’ *E*_*GABA*_ values and their total SNr input strengths, 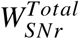 (see *Methods* for full details).

To appropriately tune the noise volatility *σ*_*η*_ and the heterogeneity factor *k*_*h*_, we constructed 400 distinct networks, with systematic variation of these parameters across networks; see Figure 2C for the collective histogram of the numbers of reciprocal SNr synaptic connections to individual neurons across the 400 slice networks. For each network, we measured the distribution of firing rates and CVs across the network and compared these distributions to those obtained experimentally. Our metric of comparison was the relative error, defined for the firing rate as 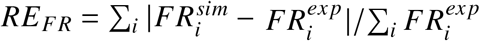, where each *FR*_*i*_ denotes the height of the probability histogram, simulated or experimental, associated with the *i*th bin in the interval of relevant firing rates (with an analogous formula for CV). We then scaled these results linearly such that the largest firing rate error was equal to 1 and the smallest to 0, and likewise for CV, to enable comparison between errors in firing rates and CVs. We found that as we varied noise volatility along with the heterogeneity parameter *k*_*h*_, better fits of the firing rate distribution and its mean were achieved for larger *k*_*h*_ and better fits of CV occurred for smaller *k*_*h*_ (Figure 2D,E,G,H). Note that a clear difference exists from left to right in each panel, with better fits to firing rate data to the right of *k*_*h*_ = 0.2 − 0.3 (Figure 2D,G) and an opposite trend for CV data (Figure 2E,H). We computed an average of the relative errors in the firing rate and CV distributions to produce an overall error score (Figure 2F) and similarly took an average of the relative errors in the means of firing rate and CV to produce an overall error score for these (Figure 2I).

We assume that a simulated SNr network matches experimental observations when a statistical comparison of the data from the two does not allow us to reject the null hypothesis that they come from the same distributions (Kolmogrov-Smirnov test, *p* > 0.05) or from distributions with the same means (t-test, *p* > 0.05). All 400 model networks were statistically different from experimental observations according to both tests both for firing rates and CVs, however. Therefore, we turned to a new network construction approach based on generating a model neuron database.

### Network Model Assembly from a Model Neuron Database Identifies Models that Match Slice Data

Although our model neuron and network connectivity parameters were carefully constrained by experimental data and we included appropriate forms of heterogeneity and noise in our model networks, their firing rates and CVs were statistically different from our recorded data. In retrospect, this disparity was not surprising, as our sweep through parameter values treated all parameters as independent, whereas some parameters may be correlated in biological neurons (*20, 21*). Thus, we adopted a new approach to leveraging our model and experimental data for network model construction.

First, we performed a new set of network simulations in which we restricted the heterogeneity to *k*_*h*_ ∈ [0.1, 0.5] and noise volatility to 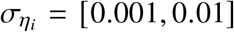, guided by Figure 2F and Figure 2I, and allowed the mean values of certain parameters related to our updates of the model relative to the previously published version (*8*), namely 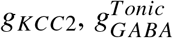, and *W*_*SNr*_ (see *Methods*), to vary. Systematic variation of these factors resulted in 1344 total 100-neuron networks, each of which we simulated.

From the outputs of these simulations, we generated a database consisting of 252 of our simulated neurons by iterating through each experimentally recorded neuron (n = 252) and identifying the model neuron from among all 134,400 simulated units (1344 simulations with 100 units each) that had the lowest error relative to that recording, defined as the average of relative error for firing rate and relative error for CV. Then, we randomly sampled 100 out of the 252 model neurons in the database, under the constraint of matching the proportions of model and experimental neurons with average firing rates in each 5 Hz firing rate bin. For example, if 25 out of 252 recorded neurons had firing rates in the 5-10 Hz bin, then we selected 10 out of our 100 model neurons from among those in the database with 5-10 Hz firing rates. We repeated this constrained, random model neuron selection for all firing rate bins; if the total number of neurons selected in this way was less than 100, either due to rounding or due to a lack of sufficiently many model neurons with firing rates in a particular bin, we randomly selected the remaining neurons from the database. All selections were made without replacement. For each set of 100 neurons selected, we constructed a model SNr network by instantiating a set of reciprocal synaptic connections among the neurons. To set the weights of these synapses, we computed a value to use as the mean for the *W*_*SNr*_ distribution based on the SNr synaptic strengths of the neurons selected from the database (see *Methods*). Similarly, we averaged the number of SNr inputs over the 100 neurons in the database and divided by 100 to compute a pairwise neuronal connection probability for the network. We repeated this assembly process with different random seeds to generate 300 tuned networks from our database of model neurons.

Out of those 300 networks, we found that 10 exhibited firing rate distributions and firing rate and CV means that statistically matched those from the recorded data (Figure 3A; Figure S1 for individual cell parameters). For example, the simulated model indicated by a green star in the lower left of Figure 3A is shown in the remaining panels of the figure. The experimental (gray) and model (green) rasters of spike times show remarkable qualitative agreement (Figure 3B), and voltage traces of model neurons from the network show a diversity of activity patterns (Figure 3C). This similarity extended to the firing rate and CV distributions and means (dashed vertical lines), considered separately, across the collection of experimentally recorded neurons and the set of model neurons in this example network, with those units that were either identified as outliers or did not fire omitted (Figure 3D,E). Moreover, the relation between firing rate and CV also showed good agreement across the experimental and computational neurons (Figure 3F). Hence, we viewed these 10 networks as successful models with which to proceed for the rest of our study.

**Figure 3:**
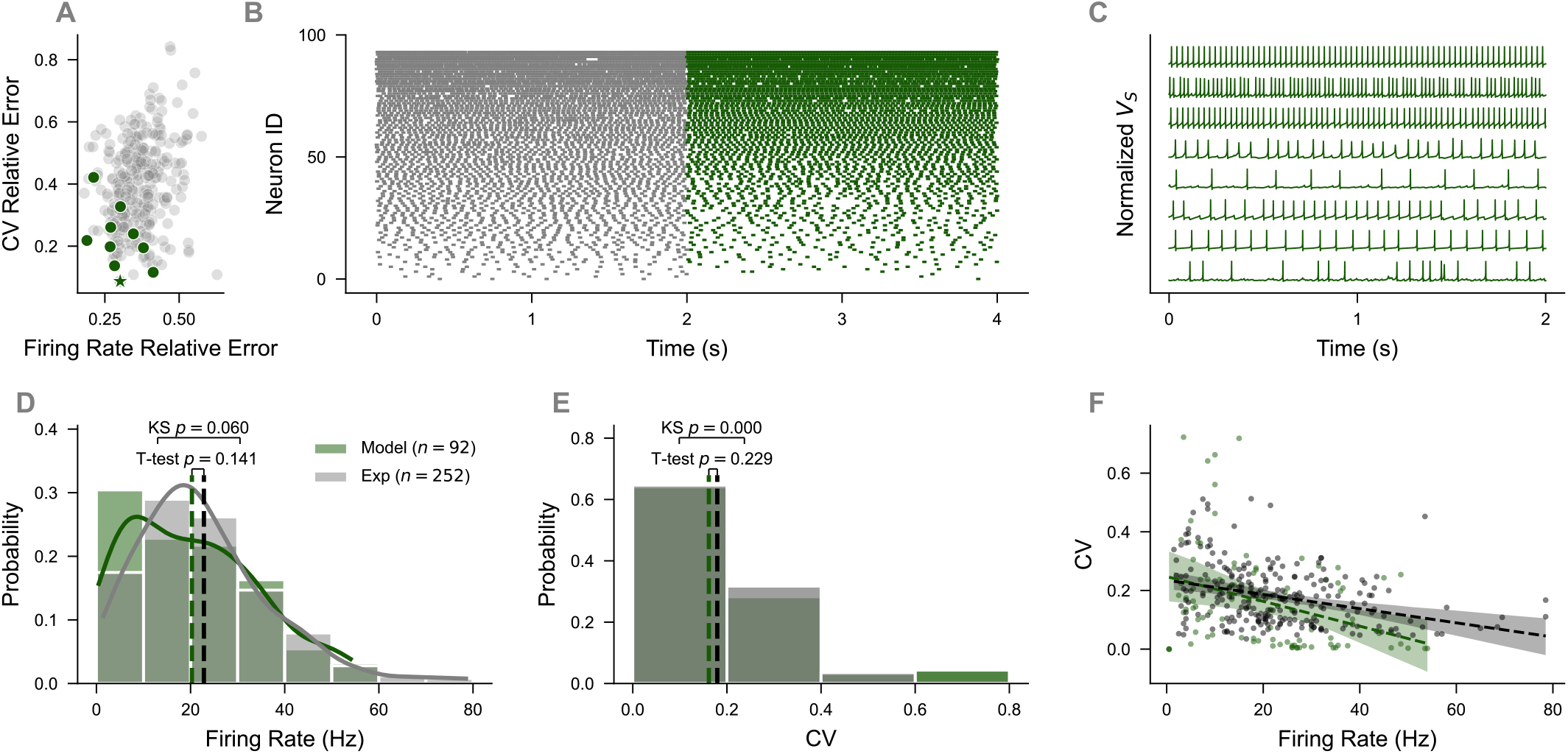
SNr slice network captures features observed in experimental recordings. (**A**) Firing rate relative error versus CV relative error based on comparison of model against experimental observations for 300 random seeds. Green points are random seeds that had non-significant p-values for K-S test and T-test for firing rate, and T-test for CV. Green star indicates the model used for the subsequent figure panels. (**B**) Raster of *n* = 92 random SNr units from experimental (model) slice recordings in gray (green). Only units with non-zero firing rate are displayed. (**C**) Voltage traces of soma compartment from 8 neurons in a model SNr network. (**D**,**E**) Panels comparing histograms of FR (**D**) and CV (**E**) for experimental data and SNr slice model. Firing rate distribution (K-S test, *p* = 0.060) and means (model *FR*_*avg*_ = 20.288 ± 13.796 Hz, exp *FR*_*avg*_ = 22.829 ± 14.208 Hz, T-test, *p* = 0.141) are not statistically different. CV distribution (K-S test, *p* < 0.001) is significantly different, while means (model *CV*_*avg*_ = 0.162 ± 0.168, exp *CV*_*avg*_ = 0.18 ± 0.1 Hz, T-test, *p* = 0.229) are not statistically different. (**F**) Linear regression with 99% confidence intervals for model (green) and slice data (gray). Model linear regression has slope = −0.004, intercept = 0.247, and *R*^2^ = 0.121 while data linear regression has slope = −0.002, intercept = 0.235, and *R*^2^ = 0.118.

### Simulated Networks Match *in vivo* Observations

An important check of the biological relevance of our SNr model networks, which were constructed initially based on slice data, was whether, with appropriate adjustments, they would also capture data recorded *in vivo* (see Figure S2A-C for a comparison of these data sets). To convert our slice model into an *in vivo* form, we assumed that all noise processes would be more prominent *in vivo*. In addition, to simulate *in vivo* conditions for SNr, we introduced inputs from the subthalamic nucleus (STN), modeled by an excitatory current with a conductance *g*_*STN*_ that evolves as an Ornstein-Uhlenbeck (OU) process over time (see *Methods* for equations). Finally, we introduced additional reciprocal connections between SNr neurons in the model network, corresponding to a doubling of the connection probability to roughly represent the transition from a slice scenario to an intact, *in vivo* setting. For each successful slice model from Figure 3A (green dots), we swept through values of the parameters associated with the OU process for 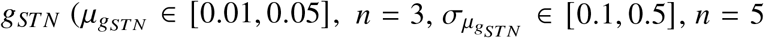, in Eq. 3 for *g*_*STN*_), as well as through 7 equally-spaced values of a scaling factor (*k* ∈ [1.1, 2.9]) for a parameter 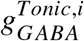 representing the intensity of ongoing GABAergic input impinging upon neurons at the soma (*i* = *S*) or dendrite (*i* = *D*); of a volatility scale factor for the 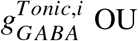 process (*k* = [2, 5, 10, 20]); and of a volatility scale factor for the current noise (*k* = [8, 20, 40, 80]; see *Methods*, in vivo *parameter search*). These parameter sweeps yielded 1,680 total *in vivo* simulations for each model slice network. With this approach, each slice model produced several *in vivo* versions that matched experimental data from *in vivo* recordings based on firing criteria (K-S test for FR and CV distributions, t-test for FR and CV means; see Figure S3 for individual cell parameters). To limit a combinatorial explosion in number of models and associated computational expense in later steps of our study, for each slice model, we kept the 3 *in vivo* models with the lowest averaged relative errors.

Figure 4A shows the scatter plot of relative errors for the FR and CV distributions of the *in vivo* models produced from the slice model in green in Figure 3A. Green points mark those simulations that were statistically indistinguishable from the experimental data, and the star point is the example model that we display in the remaining panels. The experimental (gray) and model (green) spike time rasters show strong qualitative agreement (Figure 4B), similar to that in slice. Sample voltage traces of model neurons from the *in vivo* network show a clear increase in noise and irregularity as expected in the *in vivo* setting (Figure 4C). For this network, the units had an average of 3.48 connections and a median of 3 connections from other SNr neurons (Figure 4D). Figure 4E-G are analogous to panels of Figure 3, except that here network outputs are compared against *in vivo* experimental observations. We observe that the model and data produce nearly identical firing rate and CV distributions (FR K-S test, *p* = 0.871, CV K-S test *p* = 0.811) and, especially, values of average firing rate and average CV. The joint distributions between firing rate and CV also show an excellent agreement (Figure 4E), implying that our SNr network model reproduces the relationship between firing rate and CV that arises in experimental observations.

**Figure 4:**
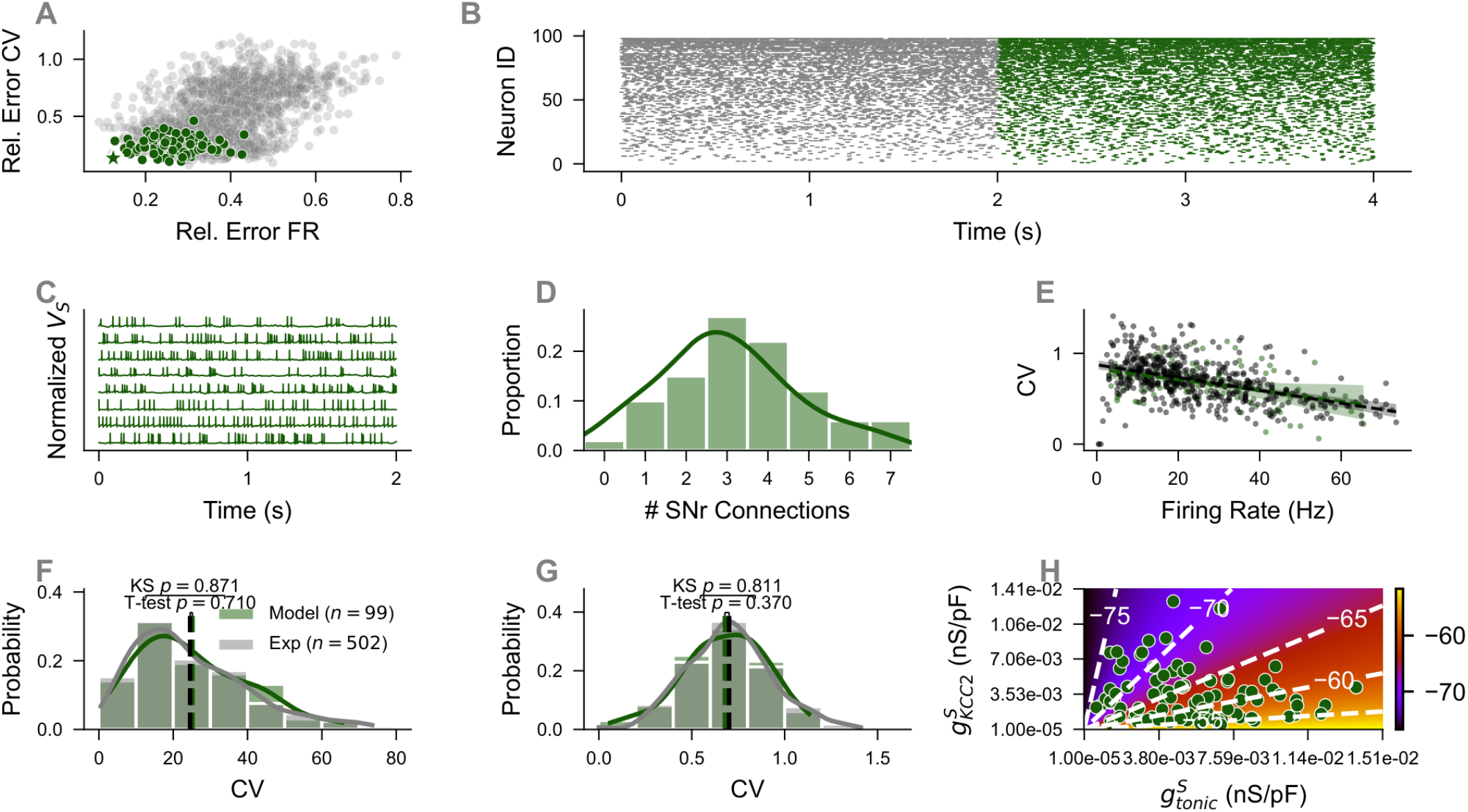
Figure depicting example *in vivo* SNr network. (**A**) All *in vivo* simulations (n=1680) from one slice seed. Simulations passing (failing) statistical criteria are green (gray) and the following panels are from the star simulation. (**B**) Raster of *n* = 99 random SNr units from experimental (model) *in vivo* recordings in gray (green). Only units with non-zero firing rate are displayed. (**C**) Voltage traces of soma compartment from 8 neurons in example *in vivo* model SNr network. (**D**) Histogram of number of SNr connections per neuron in the model network, with an average of 3.48 connections and median of 3 connections. (**E**) Linear regression with 99% confident intervals comparing model (green) and slice (gray). Model linear regression has slope = −0.006, intercept = 0.826, and *R*^2^ = 0.150 while Exp linear regression has slope = −0.007, intercept = 0.874, and *R*^2^ = 0.220. (**F**) Histogram comparing baseline experimental firing rate distribution (gray) with baseline model firing rate distribution (green). Firing rate distribution (K-S test, *p* = 0.0.871) and means (model *FR*_*avg*_ = 25.167 ± 14.588 Hz, exp *FR*_*avg*_ = 24.546 ± 15.209 Hz, T-test, *p* = 0.710) are not statistically different. (**G**) Histogram comparing baseline experimental CV distribution (gray) with baseline model CV distribution (green). CV distribution (K-S test, *p* = 0.811) and means (model *CV*_*avg*_ = 0.678 ± 0.221 Hz, exp *CV*_*avg*_ = 0.701 ± 0.229 Hz, T-test, *p* = 0.370) are not statistically different. (**H**) Heatmap for 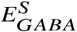 values in soma compartment for all model units in 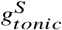 versus 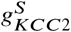 space, 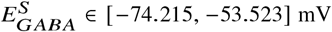 and average -63.013 mV.

### Simulations Predict Changes in Neural Properties Associated with Dopamine Depletion (DD)

Up to this point, we have discussed the development of a collection of *in vivo* SNr network models that capture baseline network firing rate and CV characteristics. We also extended these models beyond control conditions to determine what manipulations, among those proposed in past studies, best replicate SNr firing properties observed experimentally under dopamine depletion (DD).

To begin that process, we sought to consider what changes applied to a network model, which reproduced slice experiments when simulated under slice conditions and *in vivo* experiments when simulated under *in vivo* conditions, would allow it to replicate recorded SNr activity under DD (see Figure S2D-F for a comparison of control and DD *in vivo* data). To address this question, we started from each of our successful slice models (*n* = 10); for each, we selected the three descendant *in vivo* models that yielded the best match to experimental data (*n* = 30); and we generated a collection of further descendant models from these by independently sweeping through parameter values representing a variety of changes theorized to occur under DD. Specifically, experimental observations suggest that 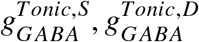 (*4, 5, 10*); 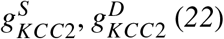; and *g*_*TRPC*3_ (*23, 24*) are all weakened under dopamine depletion whereas tonic STN activity, represented in our model by *g*_*STN*_, is increased (*4, 5, 25*). We also allowed for the possibility of increases in channel noise volatility terms, *σ*_*S*_ and *σ*_*D*_, since we reasoned that moving away from baseline, homeostatic conditions could introduce more variability into channel dynamics.

For each parameter *p* theorized to decrease under DD, we swept through multiplicative scalings *k*_*p*_ ∈ [0.33, 0.66, 1]. For each channel noise volatility parameter, we considered the multiplicative scalings 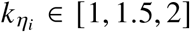, while for *g*_*STN*_ we used the scaling factors *k*_*gSTN*_ ∈ [1, 1.25, 1.5]. Note that in all cases we included *k*_*p*_ = 1 to allow for the possibility that changes in each parameter are not important for capturing DD activity. These 3 possible values for each of 7 potentially modulated parameters gave 2,187 possible parameter combinations considered for each of the 30 *in vivo* models, resulting in 65,610 total DD models. Out of these, 5,609 simulations satisfied our criteria for matching our experimentally recorded DD data (non-significant K-S test for FR and CV distributions and non-significant t-test for FR and CV means), giving a good fit rate of approximately 8.5%.

From these results, we performed several different forms of analysis to extract the dominant trends. First, over the collection of good-fitting networks and parameter sets, we computed the percentage for which each of the 7 varied parameter scaling factors *k*_*p*_ took each of its 3 possible values (Figure 5A-G). The most common scaling factor across all 7 of these parameters was 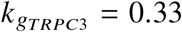, arising in approximately 64.6% of good fits, followed by 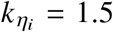, occurring in approximately 52.2% of good fits. Interestingly, three parameters, 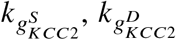, and 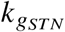, had scaling factors of 1.0, or no change, as the most common choice associated with good fits. We also extracted the 10 parameter combinations with the best coverage score (number of good fits for that parameter combination divided by number of models with that parameter combination). In Figure 5H, moving left to right, the coverage score bars for these 10 parameter combinations are each color-coded by the values of parameter scaling factors appearing in Figure 5A-G. For example, the top bar represents the parameter combination 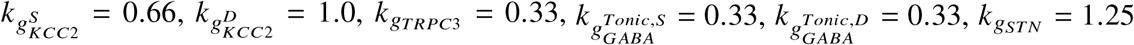, and 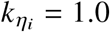, which had coverage score of 76.6% (23 good fits / 30 models). Interestingly, the other two parameter combinations that had a coverage score of 76.6% both had 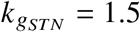 and otherwise agreed with the first one except in their 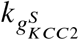 and 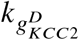 values. In all of these top 10 parameter combinations, 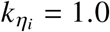, indicating that despite the prevalence of 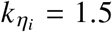 values in successful models, elevated noise volatility is unlikely to be a key feature shaping SNr activity in DD. Additionally, 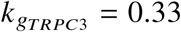 for 9/10 best models (and 0.66 in 1/10) which reinforces the finding (Figure 5C) that reductions in *g*_*TRPC*3_ are pivotal for changes in SNr activity in DD.

**Figure 5:**
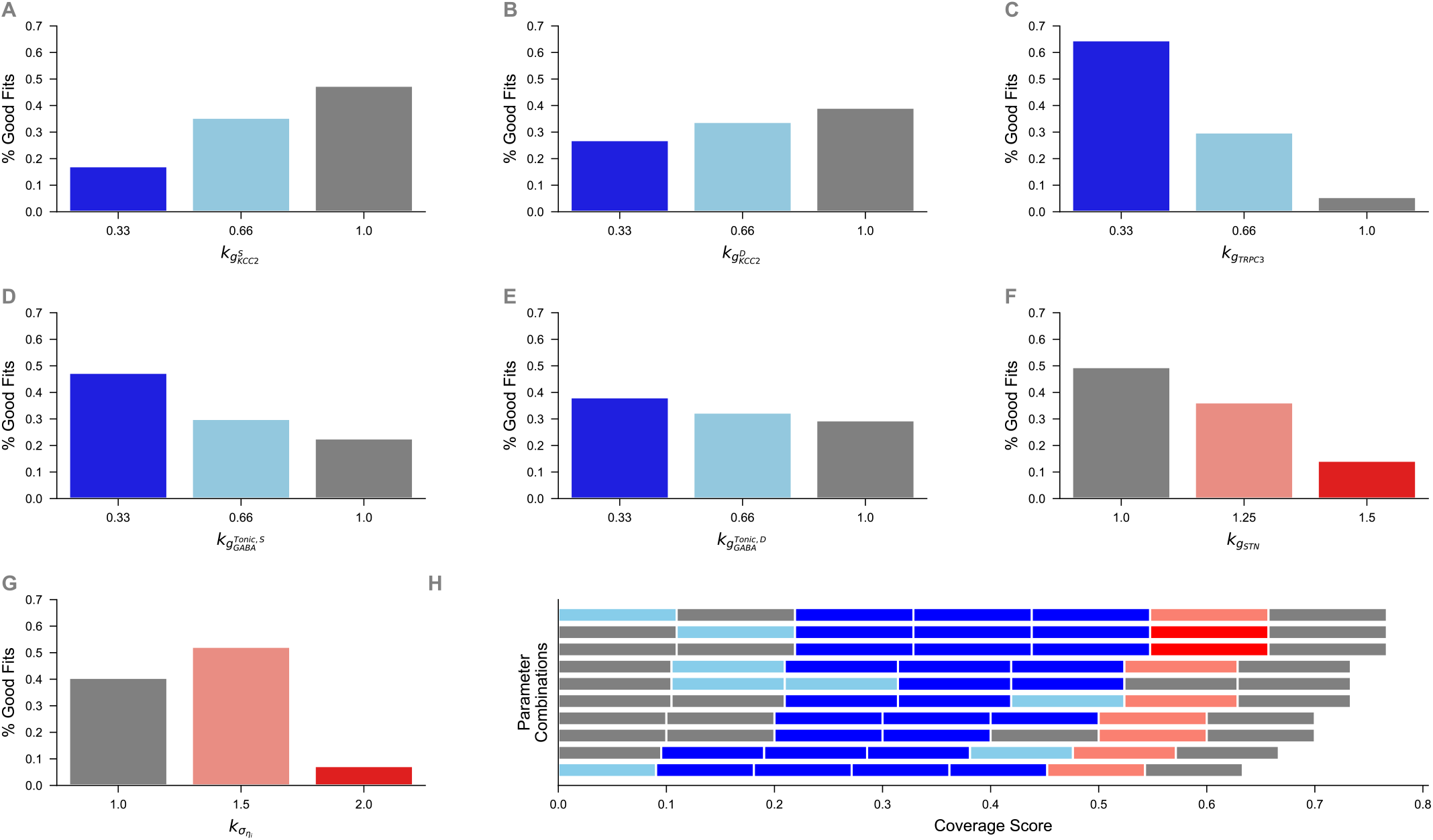
Good fit rates for parameter choices and parameter combinations. (**A**-**G**) Percentage of parameter choice (horizontal label) of all good fits (*n* = 5609), colored by decrease (blues) or increases (reds). (**H**) Proportion of *in vivo* models (*n* = 30) for most frequent successful parameter combinations (best 10 shown, from 2187 possibilities). Horizontal bars are partitioned into 7 equal length bars representing parameter choice from panel A to panel G and colored accordingly (e.g. left most patch corresponds to 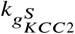, followed by 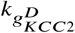, continuing until 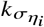).

Secondly, to analyze the hierarchy and relationships among these parameter changes more deeply, we constructed a decision tree with a depth of five (i.e., five steps beyond the starting point or root), where we kept only those paths, corresponding to successively fixing parameters at certain values, that yielded monotonically increasing conditional good fit rates (cGFRs) at each layer (Figure S4). The root starts with all parameters free and with the overall good fit rate of the DD models, 8.5%. From there, we obtain four initial branches corresponding to possible initial choices that increase the cGFFR: 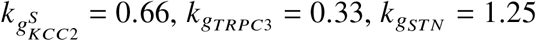, and 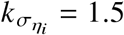. These four choices then branch into multiple paths, which all culminate at the maximum depth of five with cGFR = 44%; in fact, the nodes at depth five all represent the same 5 parameter specifications (the five listed above plus 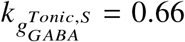), just obtained in different orders. Interestingly, by the third layer, *g*_*TRPC*3_ is the only parameter that is changed on every path, a distinction that continues at the fourth layer as well.

Our first two forms of analysis focused on specific choices of parameter values, either taken together (Figure 5) or one by one (Figure S4), that yield high probabilities of DD-like network activity. As a final approach to determining parameter importance, we constructed a decision tree based on directional changes. That is, each decision was based either on keeping a parameter at its baseline value (i.e., *k*_*p*_ = 1.0) or changing it (i.e., *k*_*p*_ < 1 or *k*_*p*_ > 1) in the direction corresponding to the parameter options indicated in Figure 5. Figure 6 shows the resulting tree, with a depth of four (i.e., four choices made per path). At the root, no choices have been made so the tree indicates an initial cGFR given by our overall simulation success fraction of 8.5%. The first decision condition provided by the tree is 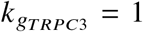. If we wish to consider what happens when this condition is false, then the left branch of the tree should be followed and analysis will be restricted to cases where *g*_*TRPC*3_ is decreased, leading to a cGFR of 12.1%. On the other hand, if we want to consider cases where the first condition holds, then we follow the initial right branch of the tree and see that when 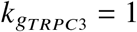, the cGFR drops to 1.4%. We can continue this process to move deeper into the tree; once each decision is made, its condition is maintained across all subsequent decisions and hence the number of models that satisfy all accumulated conditions decreases. For example, if we follow the “False” branch at each of the first two decision points, then we consider those models with 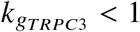 and 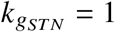 and find that among these, cGFR=17.2%. Moving through the tree, we find that there are several paths that exceed the initial cGFR = 8.5%. If we label each branch choice by either *F* for False or *T* for True, then the best performing path is *FFFT*, giving a cGFR= 23.7%, nearly three times our initial cGFR. This is closely followed by *FTTF* at cGFR= 22.2%. These two paths have, respectively, 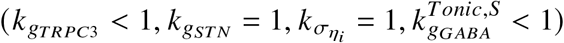 and 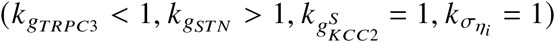. There is only one path with an initial *T* choice where the final cGFR exceeds the initial, *TFFT*, at cGFR= 11.4% while there are 7/8 leaves that exceed the initial cGFR with an initial *F* choice.

**Figure 6:**
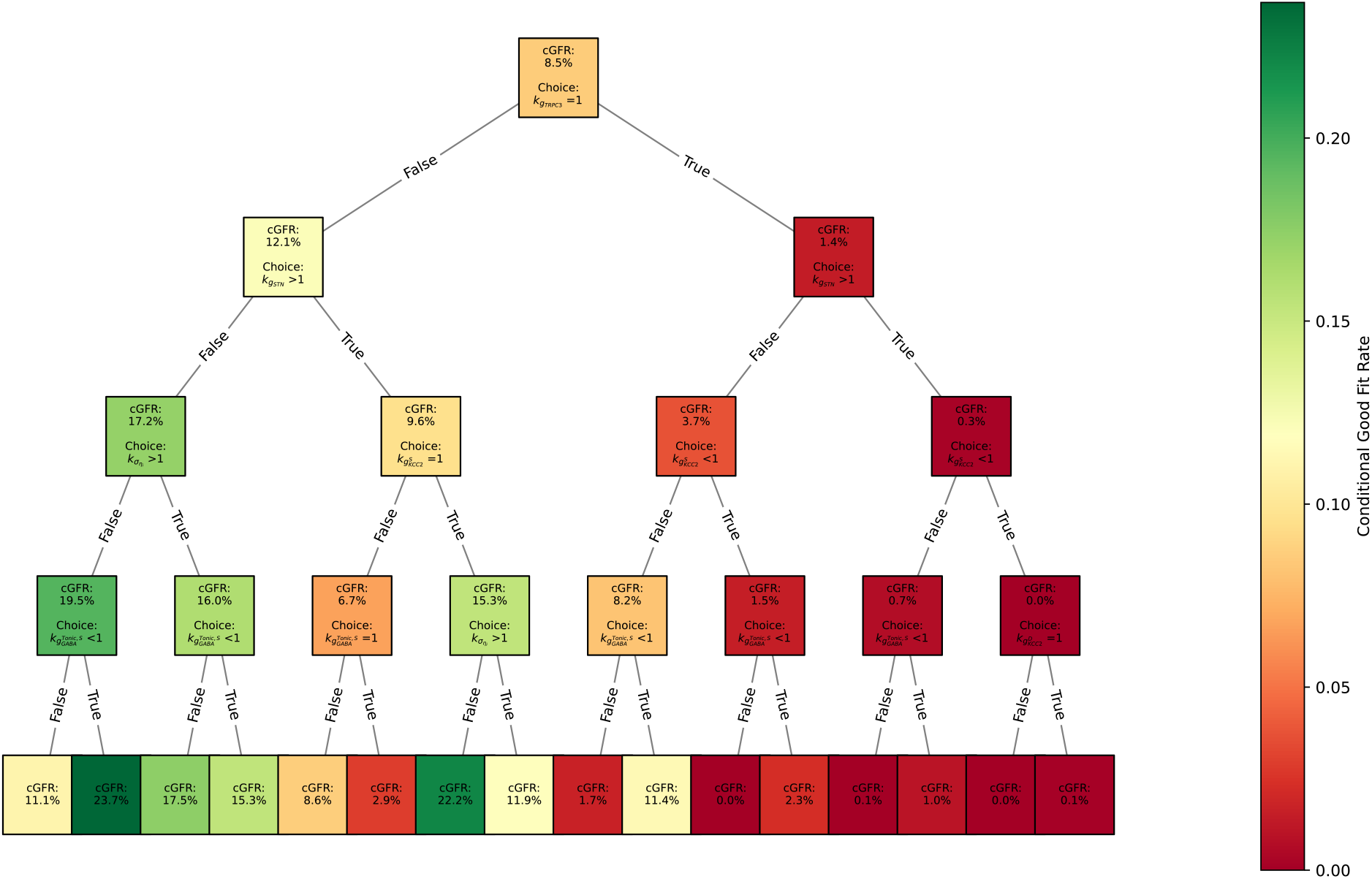
Decision tree for dopamine depleted changes. A decision tree based on changing a parameter (decrease or increase) or keeping the parameter constant (*k*_*i*_ = 1). Nodes include conditional good fit rate (cGFR) following that path and then the choice for the next branches. The top node assumes that no choices has been made giving cGFR as 8.5%, or total good fits / total dopamine depletion simulations.

Altogether, the consistent, unifying conclusion across this set of results is that a decrease in *g*_*TRPC*3_ is the key parameter change for altered SNr firing in DD. In Figure S5, we see that fixing 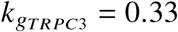, corresponding to a 2/3 reduction of *g*_*TRPC*3_ values, is enough to significantly alter model SNr firing rate and CV distributions and the relation between firing rate and CV, in similar ways to what is observed in the comparison between naive and DD experimental recordings (Figure S2, D-F).

Beyond changes in SNr firing associated with DD, another interesting aspect of SNr activity demonstrated in previous work experiments is that these neurons exhibit a diversity of responses to stimulation of the GABAergic terminals that innervate SNr from GPe and striatum (*7, 8, 10*). Thus, with our tuned network SNr model, we next studied whether the included elements are sufficient to reproduce this response diversity and whether the model provides any insights into the origins of this diversity.

### Simulated Network Models reproduce naive and DD *in vivo* SNr responses to GPe and Str stimulation

The parameter tuning in our model produced a heterogeneity of 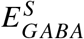 values (Figure 4F), corresponding to heterogeneity in 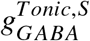, representing varying intensities of ongoing, background inhibitory inputs to different SNr neurons, and 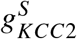; the result for 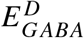 in the dendrite is not shown but is analogous. Based on past results (*8*), we expect the distribution of 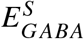 values, with an average of -63.013 mV and a range from -74.215 mV to -53.523 mV, to translate into a diversity of response types under simulated stimulation of the inhibitory synaptic afferents to the network.

### GPe Stimulation

Experimental recordings were made in SNr neurons during ten trials each consisting of one second of 20Hz stimulation of GPe terminals in SNr (found previously to give similar results to other stimulation rates between 10 and 60 Hz (*8*)). We simulated these experiments in model SNr networks by first simulating each network under 5 seconds of baseline, unstimulated conditions followed by 10 stimulation trials of 3 seconds each, for a total of 35 simulated seconds, akin to the experimental procedure detailed in (*10*) (for more details see *Methods*). Each stimulation trial consisted of one second of baseline simulation, which we call the *pre-stim* period, followed by 20Hz pulses, each lasting for one time step (*dt*), for one second of simulated time, followed by a one second *post-stim* period after the end of stimulation. The governing equations for the stimulation-induced input to the model SNr neurons were left unchanged from an earlier computational study (*8*). Recent work has shown correlated variation in the strength of GPe synaptic inputs to SNr neurons, *W*_*GPe*_, and the decay time constant, 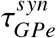, of these inputs (*10*). We used that data to generate a linear regression of 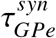 versus *W*_*GPe*_ values. Then, we selected *W*_*GPe*_ values uniformly at random from the experimentally derived *W*_*GPe*_ range and for each, we randomly selected 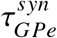 from a 99% confidence interval around the linear regression line at that *W*_*GPe*_. Finally, to fit the data, we scaled all *W*_*GPe*_ values by a multiplicative factor of 1/10 (see *Methods*). For these simulations, as in the DD case, we used the three *in vivo* models with lowest error scores produced from each of the 10 slice models (Figure 3A) and ran three full, 35-second simulations of each, with a different random seed each time; thus, we considered 90 different 100-neuron model runs altogether.

We found that our model networks nicely recapitulated the SNr responses observed experimentally (*10*). We initially quantified these responses using a modulation factor, 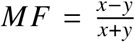, where *x* represents the firing rate during the stimulus period and *y* represents the firing rate during the pre-stim period, and observed an excellent agreement between the MF distributions from the experiments and simulations (Figure 7A,B; 90 different 100-neuron models, resulting in 9000 MFs; experimental cells: average modulation factor *MF*_*exp*_ = −0.033 ± 0.131, model cells: average modulation factor *MF*_*model*_ = −0.015 ± 0.102, not statistically different via a t-test, *p* = 0.109; MF distributions not statistically different via a K-S test, *p* = 0.638). We used an objective algorithm encoded in software that is freely available online (STReaC, https://github.com/jparker25/streac, (*9*)) to classify each recorded and each simulated neuron’s response to stimulation, relative to its prestim activity, into one of five categories: no effect, complete inhibition, partial inhibition, adapting inhibition, or excitation. We observed that the predominant outcome of GPe stimulation in experiments and simulations was no effect on SNr firing (Figure 7C, grey; note that y-axis starts at 80%), while the overall distribution of response types across the two conditions were not statistically different (χ^2^ test, p=0.168) and, when compared individually, only 3 of our 90 models differed significantly from experiments. Finally, we note that SNr response to GPe stimulation was not dictated simply by the strength and time constant (duration) of GPe input, as those responding model neurons had values that were widely scattered throughout the 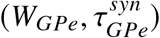 distribution (Figure 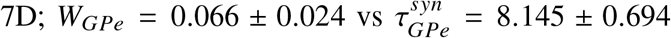, points colored by the STReaC response classification; Figure S6A for quantified response gradient).

**Figure 7:**
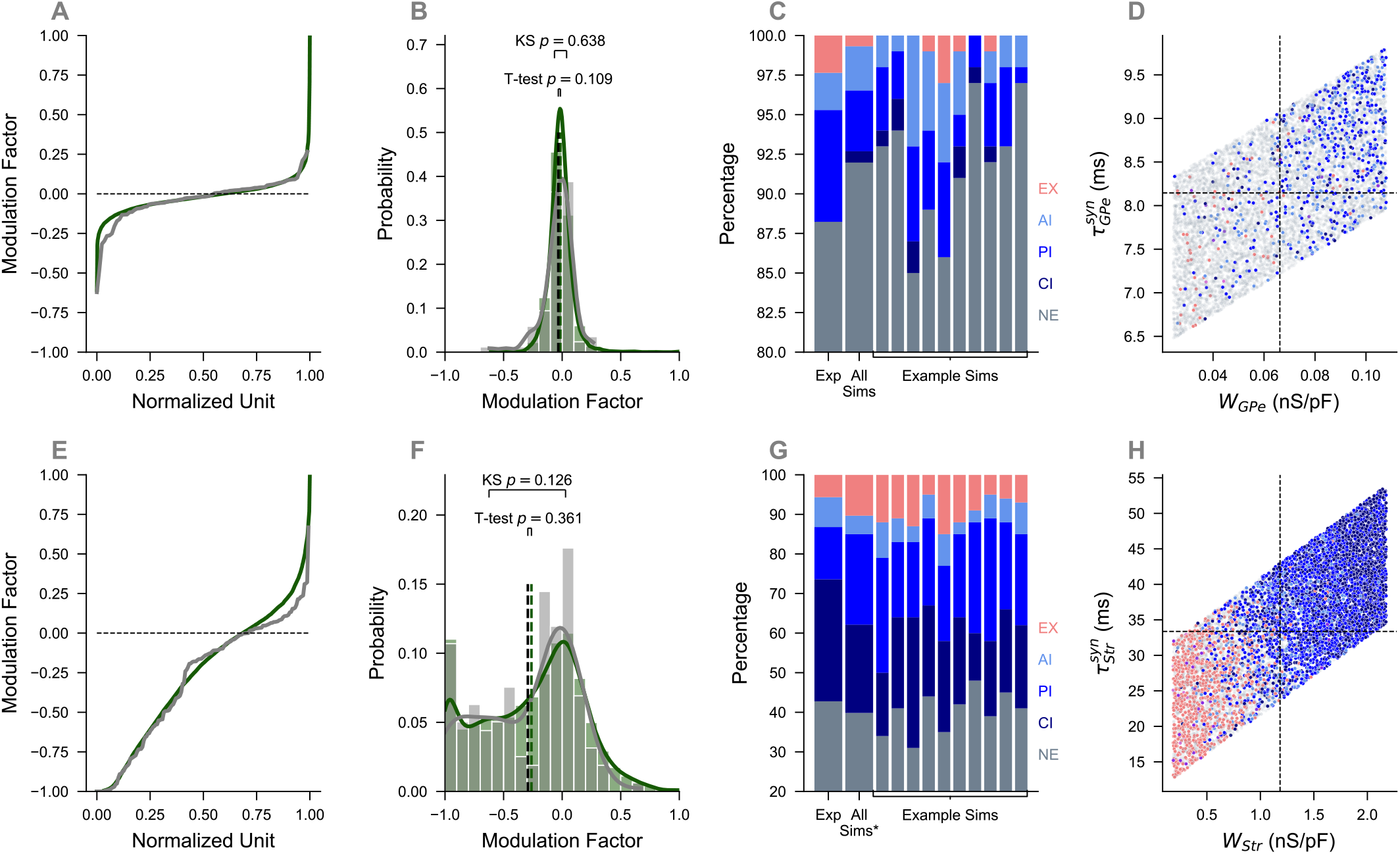
Figure depicting naive SNr model responses to stimulation. (**A**) Normalized cell number versus modulation factor for experimental responses (black) and model responses (green) to GPe stimulation. (**B**) Histogram of modulation factors from panel (**A**). Distributions (K-S test, *p* = 0.638) and means (*MF*_*model*_ = −0.015 ± 0.102, *MF*_*exp*_ = −0.033 ± 0.131, T-test, *p* = 0.109) statistically agree. (**C**) Experimental and simulated networks STReaC response distribution to GPe stimulation. All GPe sims collected together were not statistically significant (χ^2^, *p* = 0.168). Only 3/90 sims were statistically significant. Ten randomly selected examples are shown. (**D**) *W*_*GPe*_ (nS/pF) (*W*_*GPe*_ = 0.066 ± 0.024 nS/pF) connection strengths versus 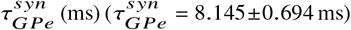 decay constants. Colored by responses as defined by STReaC toolbox. (**E**) Normalized cell number versus modulation factor for experimental responses (black) and model responses (green) to D1 stimulation. (**F**) Histogram of modulation factors from panel (**E**). Distributions (K-S test, *p* = 0.126) do not statistically agree and means (*MF*_*model*_ = −0.262 ± 0.425, *MF*_*exp*_ = −0.293±0.396, T-test, *p* = 0.361) do not statistically agree. (**G**) Experimental and simulated networks STReaC response distribution to Str stimulation. All Str sims collected together were statistically significant (χ^2^, *p* = 0.001) due to 50/90 sims were statistically significant. Ten randomly selected examples are shown. (**H**) *W*_*Str*_ (nS/pF) (*W*_*Str*_ = 1.184 ± 0.582 nS/pF) connection strengths versus 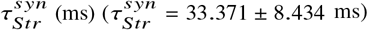 decay constants. Colored by responses as defined by STReaC toolbox.

### Striatal D1 SPN Stimulation

For simulations of stimulation of D1 SPN terminals in SNr, we used the same stimulation protocol as with GPe stimulation, and values of input decay constants, 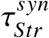, and strengths, *W*_*Str*_, were determined from experimental data (*10*) following the same procedure as was used for GPe inputs. One additional modeling aspect in this case was that we introduced anti-correlations between the *W*_*Str*_ values assigned to model SNr neurons and their dendritic GABA reversal potentials, 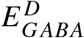, which improved the model fit to the data, especially in terms of agreement of modulation factors, and hence represents a prediction of this modeling work (Figure S7).

Similarly to the case with GPe stimulation, the simulated and recorded SNr responses to D1 stimulation yielded strong agreement in modulation factor distributions (Figure 7E-F; experiments: average modulation factor *MF*_*exp*_ = −0.293 ± 0.396, simulations: average modulation factor *MF*_*model*_ = −0.262 ± 0.425, means not statistically different via a t-test (*p* = 0.361), distributions not statistically different via a K-S test (*p* = 0.126)). The STReaC classification of SNr responses showed qualitatively similar response distributions when comparing the aggregate results from all experiments to the aggregate from all simulations as well as to individual simulations (Figure 7G), although there were statistically significant differences between the aggregates based on a χ^2^ test (*p* < 0.01) and between the experimental results and 50 out of 90 individual models. Although these statistical tests show there are response differences, we note that the MF analysis and STReaC results are nonetheless similar qualitatively across experimental and simulated results.

Finally, with D1 stimulation, SNr response type was much more linked to the strength and time constant of the input than in the GPe case. Labeling of points in the 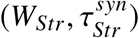 distribution based on their response (Figure 7H, *W*_*Str*_ = 1.184 ±0.582 nS/pF, 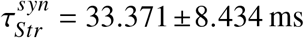) reveals a clear gradient from *complete inhibition* responses (dark blue) to *excitation* responses (light pink) as *W*_*Str*_ and 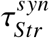 decrease (Figure S6B for quantified response gradient). Hence, we infer that relatively weak and short-lived inhibition from striatum yields a high likelihood of an excitatory response, but is not necessary for such a response to occur, based on the scattering of pink dots at larger values in the distribution.

Overall, we see that SNr responses to GPe stimulation are often weak, sometimes inhibitory, and occasionally but rarely excitatory, and our simulations predict that which cells are inhibited is not simply dictated by the strength or duration of the GPe inputs that they receive. In contrast, SNr neurons have a higher probability of responding to stimulation of terminals from D1-expressing SPN neurons in striatum. These responses can be inhibitory or excitatory, and which case occurs is strongly linked to the strength and duration of the inputs that the target SNr neuron receives from the D1-SPNs. Before diving deeper into the factors shaping SNr responses to stimulation, we considered changes in SNr responses to stimulation under DD.

### GPe Stimulation under DD

Similarly to our approach for the naive SNr model network, we simulated the experimental stimulation protocol in the dopamine depleted SNr model networks. Example results for GPe stimulation under dopamine depleted conditions using the 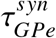 and *W*_*GPe*_ tuned values are shown in Figure 8A-D and are analogous to the naive case (Figure 7A-D).

**Figure 8:**
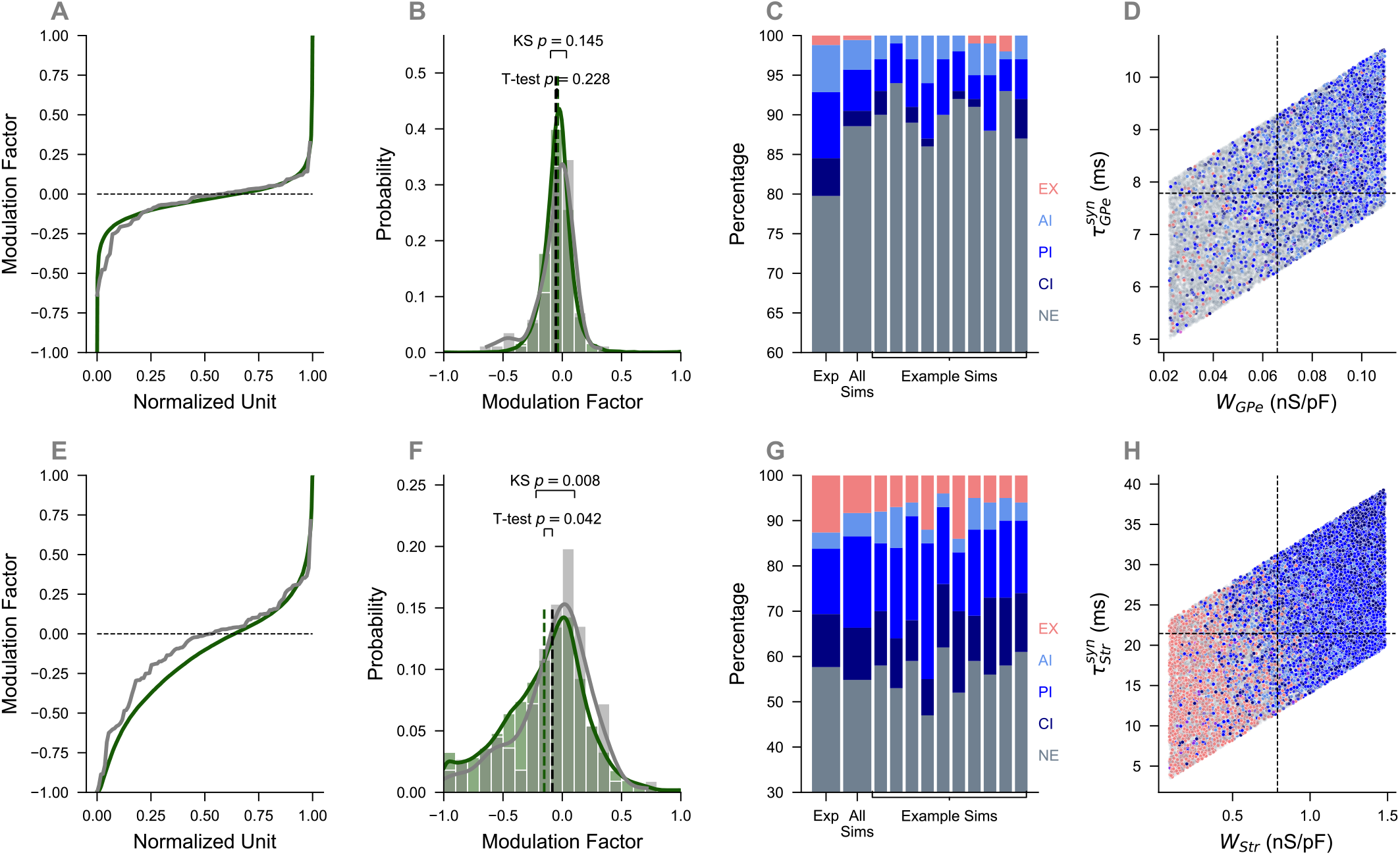
Figure depicting DD SNr model responses to stimulation. (**A**) Normalized cell number versus modulation factor for experimental responses (black) and model responses (green) to GPe stimulation. (**B**) Histogram of modulation factors from panel (**A**). Distributions (K-S test, *p* = 0.145) do not statistically agree and means (*MF*_*model*_ = −0.038 ± 0.128, *MF*_*ex p*_ = −0.054 ± 0.156, T-test, *p* = 0.228) do statistically agree. (**C**) Experimental and simulated networks STReaC response distribution to GPe stimulation. All GPe sims collected together were not statistically significant (χ^2^, *p* = 0.115). Only 25/270 sims were statistically significant. Ten randomly selected examples are shown. (**D**) *W*_*GPe*_ (nS/pF) (*W*_*GPe*_ = 0.066 ± 0.025 nS/pF) connection strengths versus 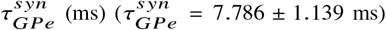 decay constants. Colored by responses as defined by STReaC toolbox. (**E**) Normalized cell number versus modulation factor for experimental responses (black) and model responses (green) to D1 stimulation. (**F**) Histogram of modulation factors from panel (**E**). Distributions (K-S test, *p* = 0.008) and means (*MF*_*model*_ = −0.152 ± 0.353, *MF*_*ex p*_ = −0.084 ± 0.315, T-test, *p* < 0.042) do not statistically agree. (**G**) Experimental and simulated networks STReaC response distribution to Str stimulation. All Str sims collected together were not statistically significant (χ^2^, *p* = 0.292). Only 21/270 sims were statistically significant. Ten randomly selected examples are shown. (**H**) *W*_*Str*_ (nS/pF) (*W*_*Str*_ = 0.790 ± 0.401 nS/pF) connection strengths versus 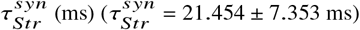 decay constants. Colored by responses as defined by STReaC toolbox.

The sorted MF values for the 270 different models (27,000 total MFs) shown in green and the corresponding experimental values plotted in black exhibit a substantial overlap (Figure 8A). A comparison of the MF distributions across simulations and experiments shows an average modulation factor of *MF*_*exp*_ = −0.054 ± 0.156 for the experimental cells and and an average modulation factor of *MF*_*model*_ = −0.038 ± 0.128 for the model cells (Figure 8B). These means are not statistically different based on a t-test (*p* = 0.228), nor are the distributions statistically different based on a K-S test (*p* = 0.145), implying that, as in the naive case, the DD model cells’ firing rate changes in response to simulated GPe stimulation match the responses of experimental cells under DD conditions. Naturally, the STReaC response distributions showed some variability across simulated neurons. The combined response proportions from all of these distributions were not statistically different from the experimental response distribution (Figure 8C, χ^2^ test, *p* = 0.115) and only 25/270 individual models exhibited statistically significant differences from experimentally recorded response distributions. The distribution of 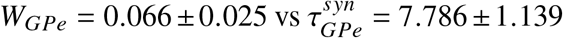 (ms) colored by the STReaC response (Figure 8D) reveals that the vast majority of the cells are gray, similar to the naive model, indicating a *no effect* response. Compared to the naive case (Figure 7D), the responsive cells generally lay in the upper right quadrant of the GPe synaptic parameter space (Figure S6C for quantified response gradient).

### D1 Stimulation under DD

Simulation of D1 stimulation in the same 270 DD models reveals an approximate agreement between the experimental modulation factors (black) and the simulated modulation factors (green) (Figure 8E-H). More precisely, the curves agree closely with *MF* > 0 but have a clear separation with *MF* < 0 (Figure 8E). A comparison of the histograms of these two sets of values shows that the average modulation factor of the experimental cells (*MF*_*exp*_ = −0.084 ± 0.315) does differ statistically from the average modulation factor of the model cells (*MF*_*model*_ = −0.152 ± 0.353) based on a t-test (*p* = 0.042). The full distributions are also statistically different via a K-S test (*p* < 0.01), yet we still find that the STReaC responses are qualitatively quite similar between experiments and simulations (Figure 8F-G). Moreover, the full set of experimental responses and the collection of all simulated responses were not significantly different (χ^2^ test, *p* = 0.292) and only 21/270 individual simulations exhibited a statistically significant difference from the experimental responses (Figure 8G). Again, we note that the MF analysis and STReaC results are mutually complementary and agree qualitatively with the experimental results. We found a similar gradient of STReaC responses in the parameter space of 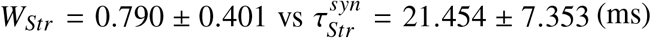 values (Figure 8H; Figure S6D for quantified response gradient) as in the analogous naive case (Figure 7H).

### Response Class Mechanisms

Given the variety of statistical and qualitative agreements between the responses to stimulation of inhibitory terminals onto SNr neurons in our model network simulations and those observed experimentally, we sought to use additional analysis of the factors shaping the model responses to provide predictions about the biological mechanisms involved. Specifically, across all neurons showing a certain type of response to GPe input, we examined the GABA reversal potential 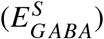, the synaptic decay time constant 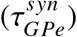, the synaptic input strength from GPe (*W*_*GPe*_), the strength of reciprocal synaptic input from other SNr neurons 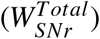, and the modulation factor *for those SNr neurons projecting to a given neuron (MF*_*input*_), separately for the naive and dopamine depleted conditions. Similarly, for responses to Str input, we considered 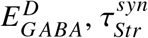, and *W*_*Str*_, along with 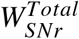 and *MF*_*input*_. Note that *MF*_*input*_ is dimensionless; values less than zero indicate a decrease in the average presynaptic firing rate and values greater than zero indicate a strengthening of this rate under GPe or Str terminal stimulation, relative to baseline conditions.

Using the response classification from STReaC (Figure 7 and Figure 8) we grouped the responses into three primary groups: no effect, inhibition, and excitation. Then, we determined the distribution of parameter values for the inhibition and excitation groups in each condition (GPe or Str stimulation, naive or DD setting) to determine if any are indicative of the response class, based on comparison to the no effect distribution. To ensure a fair comparison, we normalized the no effect distribution such that the mean of the distribution was zero and the standard deviation was one, normalized each of the inhibition and excitation distributions to the values of the no effect distribution, and then determined the Wasserstein distance (WD) based on these normalized distributions. The WD, also known as the “earth-mover’s distance” measures how much “work” is needed to equalize two distributions; that is, it computes how much probability mass must be moved to transform one of the distributions into the other. Larger WD values indicate a larger difference in the normalized response distribution compared to the normalized no effect distribution, compared to smaller WD values. Since the WD value is not directional, we additionally determine the direction of the difference by comparing the means of the distributions (Figure S8).

In the naive case, we found that 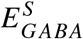 and 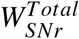 were the primary factors underlying inhibited firing responses to GPe stimulation and *MF*_*input*_ was the least relevant factor (Figure 9A). Similar results held for inhibited responses to striatal stimulation (with 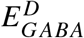 in place of 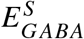), with strength of striatal inputs (*W*_*Str*_) also playing an important role (Figure 9C). Interestingly, WD values associated with striatal stimulation were larger than the WD values for GPe stimulation, very roughly by a factor of two.

**Figure 9:**
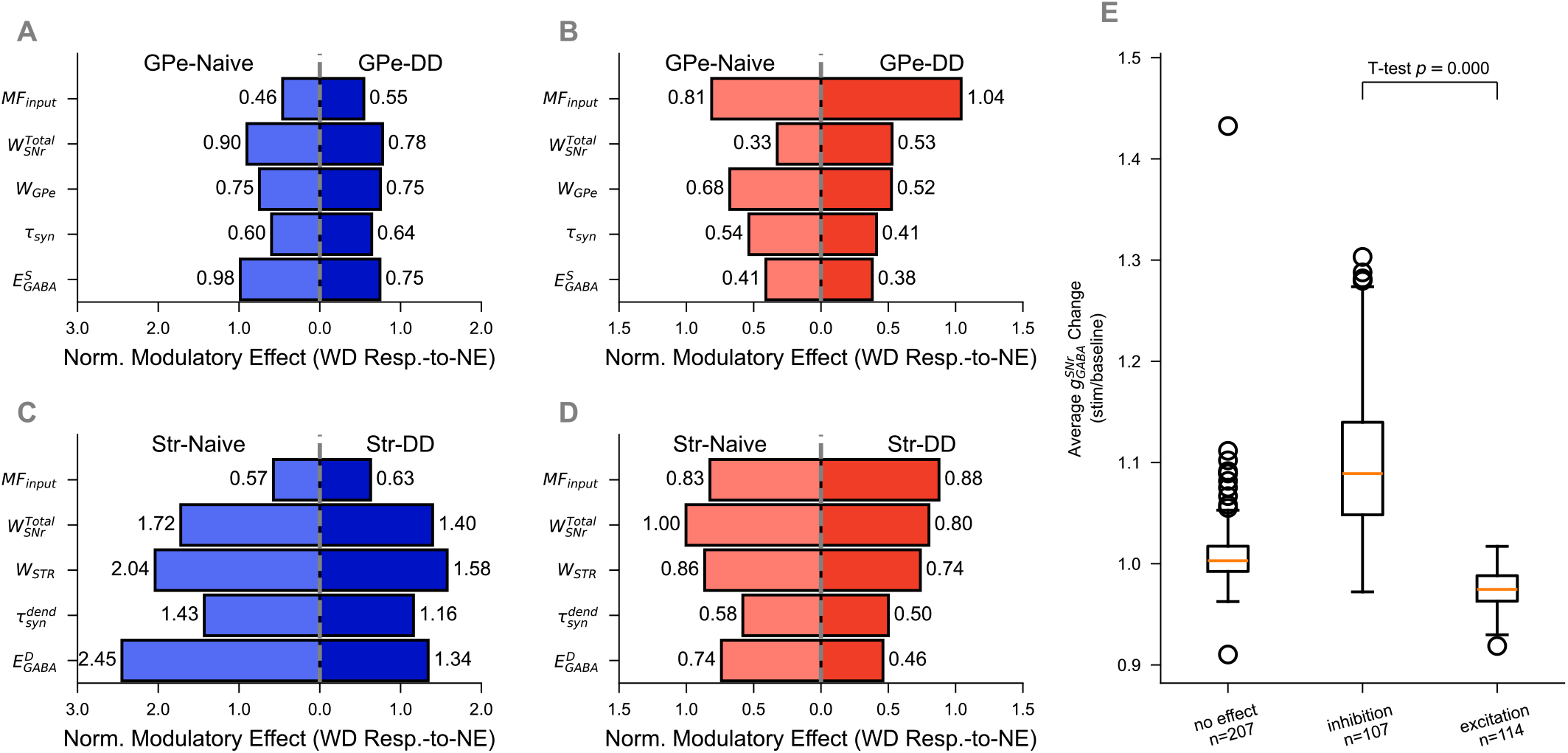
Wasserstein distance (WD) for response class compared to no effect distributions. (**A-B**) WD values for SNr (**A**) inhibitory responses and (**B**) excitatory responses to GPe stimulation, compared to no effect distributions, for each parameter. (**C-D**) WD values for SNr (**A**) inhibitory responses and (**B**) excitatory responses to Str stimulation, compared to no effect distributions, for each parameter. Light (dark) colors represent naive (dopamine depleted) conditions. (**E**) Ratio of average 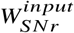 during stimulation to average 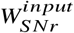 during baseline, averaged over 10 trials, grouped by response class (*N* = 5 models). Each ratio is significantly different from the other two (no effect: mean=1.01, median=1.00, *n* = 207; inhibition: mean=1.10, median=1.09, *n* = 243; excitation: mean=0.97, median=0.97, *n* = 50; NE v. Exc: *t*-test, *p* < 0.001; Exc v. Inh: *t*-test, *p* < 0.001; NE v. Inh: *t*-test, *p* < 0.001).

For excitatory responses to GPe stimulation in the naive case, the largest WD value was associated with *MF*_*input*_, followed by the strength of the input from the stimulated terminals (*W*_*GPe*_), and the smallest arose for 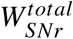 (Figure 9B). For striatal stimulation in the naive case, in contrast, the largest WD value was 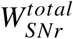, with *MF*_*input*_ and stimulated terminal input strength (now *W*_*Str*_) again large (Figure 9D). We compared the ratio of the average *W*_*SNr*_ during stimulation to the average *W*_*SNr*_ during the baseline period, averaged over trials (Figure 9E) for five naive networks. We found that for neurons with an excitatory response, the average *W*_*SNr*_ ratio was significantly lower during stimulation than for neurons that exhibited an inhibitory response (excitation: 0.975, *n* = 50; inhibition: 1.10, *n* = 243; *t*-test, *p* < 0.001) or showed no effect from these inputs (excitation: 0.975, *n* = 50; no effect: 1.01, *n* = 207; *t*-test, *p* < 0.001).

The relative contributions of the studied factors to responses occurring under DD generally remained similar to those observed in the naive case. DD diminished the majority of the WD values for both inhibitory and excitatory responses, suggesting that responses to terminal stimulation were at least slightly less predictable from the characteristics of neurons and their inputs in the DD than in the naive case, although *MF*_*input*_ was universally an exception. These decreases were most prominent for inhibitory responses to striatal stimulation (Figure 9C). Overall, the same parameters remained dominant in shaping SNr responses to GPe and Str terminal stimulation in DD as in the naive case, although relative magnitudes of the WD values changes between the two conditions; for example, in the case of Str stimulation, *MF*_*input*_ became the dominant factor for excitatory responses.

In summary, this analysis predicts that the most prominent mechanism for producing excitatory SNr responses to GABAergic terminal stimulation is generally a drop in the inhibition that an SNr neuron receives from other SNr neurons, as measured by *MF*_*input*_, presumably because they are inhibited by the stimulation. Inhibitory SNr responses, on the other hand, depend on the strength of inhibitory inputs to the target neuron both from the stimulated site and from within the SNr along with the GABA reversal potential at the compartment where the stimulated synapse was located, which was more negative on average compared to the analogous GABA reversal potential from excitatory responses (Figure S8A and Figure S8F), with less of a contribution from the timescale of the inhibition and the degree of inhibition of the SNr neurons that project to the target neuron. Under DD conditions, the WD values for the leading factors causing inhibitory responses to striatal stimulation diminished, suggesting a general drop in the predictability of which SNr neurons would show these types of responses.

## Discussion

### Model Construction

In this work, we took on a major modeling challenge: we sought to expand the single-neuron SNr model presented in (*8*) to a network-level framework capable of replicating slice and *in vivo* experimental results across baseline and stimulated conditions in naive and DD preparations. Standard approaches to model neuronal network construction form networks using model neurons with parameters that are identical or are selected randomly from distributions, typically based on either an experimentally-derived mean or range. Values of different parameters are sampled independently to derive each model neuron, and these selections are made independently across model neurons. However, we found that this approach did not yield model networks that matched experimental data (cf. Figure 2), even though we included channel noise as well as synaptic connection probabilities and strengths within the network that were tuned to match published results. Indeed, a potentially important shortcoming of this approach is that it neglects correlations among the conductance values and other properties that arise within neurons (*20, 21*). We therefore introduced a novel network assembly method in which we matched individual model neurons one-to-one with experimentally recorded neurons and then constructed model networks by sampling the resulting model neuron database. Importantly, although the experimental data consisted of single-neuron recordings, neurons in slice and *in vivo* preparations belong to networks; correspondingly, we derived our surrogate model neurons from a large collection of preliminary model network simulations. This procedure allowed for quick identification of neurons with appropriate firing properties in the presence of network effects, whereas individual model neurons may not retain firing properties when included in a network.

Even though the selected neurons were extracted from different network simulations, when we coupled the samples together into a model network, taking into account input properties present in the original simulations, a subset of the resulting networks accurately captured the firing rate and CV distributions found experimentally in slice preparations (Figure 3). Overall, a small percentage of our initial parameter samples and subsequently constructed networks fit the recorded SNr firing rates and CVs. Indeed, since we used a biologically detailed model, with components constrained by earlier experiments and known to be relevant for SNr neurons, and the match to data was not a design element of the model but rather emerged from the model dynamics, a low success rate in the fitting procedure is not surprising and reassures us that there is something special and hence informative about the successful models.

From the original successful set of networks, the introduction of factors such as additional input sources present *in vivo* yielded a collection of model networks – each with a distinct set of neurons and parameter values – that not only fit *in vivo* experimental data from naive conditions but also matched recordings from DD conditions and from optogenetic stimulation of GPe or striatal synaptic terminals in SNr in both cases. Overall, we propose that this iterative process – using a large set of preliminary, experimentally-constrained network simulations to generate a collection of candidate model neurons, assembling a database of model neurons one-to-one matched to neurons recorded from within experimental preparations, and then sampling and coupling together neurons from the database – represents a novel innovation for constructing model networks that reproduce biological network activity. Interestingly, another innovative approach has recently been applied to generate an SNr network model (*26*). That work presents a volumetric reconstruction of the SNr, with great emphasis on spatial anatomical details and with multi-compartmental model SNr neurons; our study complements that approach, with simpler individual neuron models that allow for an emphasis on the roles of individual conductances, reversal potentials, and other dynamics-related parameters and with matching of individual SNr recordings that ensures inclusion even of less common SNr neuron types.

### Changes under Dopamine Depletion

A major motivation for our model network construction was to test ideas and generate predictions about what changes in SNr neuron properties may underlie their altered activity in DD conditions.

We investigated SNr activity in DD conditions in our simulations by performing a parameter sweep on each of the *in vivo* network models, such that parameters previously predicted to change under DD (*4, 5, 10, 22–25*) were each systematically adjusted over a range of values. This approach allowed us to test the success of models with a wide variety of combinations of these parameter changes in capturing experimental observations collected *in vivo* under DD conditions. We analyzed our results in multiple ways, based on the likelihood of good fits emerging under individual parameter changes, marginalized over all values taken by other parameters, and under various combinations of parameter changes. Our results universally suggested that the most important parameter changes underlying altered activity in recorded neurons under DD were reductions in *g*_*TRPC*3_, the conductance of the TRPC3 channels found in SNr dendrites (*23, 27*) (Figures 5, 6), with larger reductions (down to 33% of original values) yielding higher likelihoods of good fits than smaller reductions (66%). Experiments have shown that TRPC3 channels are present and constitutively active in SNr GABAergic neurons and that their pharmocologic blockade reduces firing frequency and increases its irregularity (*23, 27*). Further results suggest that TRPC3 channel activation is linked to engagement of dopamine receptors, possibly via dopamine-related stimulation of lipid signaling molecules that boost TRPC3 channel activity (*28,29*). If, as these findings suggest, TRPC3 channel activation mediates dopamine-induced increases in SNr neuron excitability, then the loss of dopamine could significantly impact SNr firing (*27*), as our model predicts. Along with TRPC3 effects, even better agreement with data resulted from changes in other parameters exposed by our analysis in conjunction with decreases in *g*_*TRPC*3_. Examining specific parameter combinations that almost always matched data (Figure 5H), the parameter selections that were always present along with reductions in *g*_*TRPC*3_ were large decreases in the background GABAergic input coming to the soma of model neurons and a lack of change in the level of channel noise present. The former effect could be caused by a reduction in GPe firing that has been observed in DD (*30–32*); interestingly, this modulation could also change the GABA reversal potential and the impact of GPe on SNr acitvity under DD conditions (*8*). The idea that SNr neuron voltage dynamics could exhibit increased fluctuations under DD is consistent with reports that dopamine increases the signal-to-noise ratio in PFC neurons (*33, 34*) yet our results argue against an increase in fluctuations that would have a meaningful impact on activity patterns. Increases in conductance of STN inputs, which were modeled as a stochastically generated excitatory spike train, also were usually present in successful parameter sets, consistent with the increases in STN firing under DD predicted by classical models (*4, 5*).

Examining groups of parameter changes that jointly contributed to model fit to DD data did not add any definitive insights to these conclusions but did highlight that certain changes that were not effective on their own could yield much higher rates of good fit when implemented together. For example, when taken together with reductions in TRPC3 conductance and background GABAergic input to the soma, increases in STN input efficacy, reductions in somatic KCC2 channel conductance, and increases in channel noise could yield such a high rate. Whether such specific combinations are biologically relevant, however, remains for future exploration.

### Response Modulation

Classical models treat the BG output centers, such as the SNr in rodents, as targets of inhibitory inputs that come from the direct pathway via striatum and from the indirect pathway via the GPe (*4, 5*). Yet, as seen both in previous work (*7–10*) and in our results (Figure 7 and Figure 8), the reality is that stimulation of indirect and direct pathway terminals in SNr, consistently across a variety of frequencies, results in heterogeneous SNr responses. In the naive setting (Figure 7H), visual inspection suggests that there is a gradient in the *W*_*Str*_ and 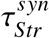 space, where stronger, longer inputs from striatum are linked with inhibitory responses in SNr; to a weaker extent, similar trends hold with GPe inputs as well, although in most cases these are classified as having no effect Figure S6B). To be clear, this designation does not mean that SNr firing after terminal stimulation remains identical to that before stimulation. Our SNr response classification was done objectively, using a published, automated classifier tool specifically developed to distinguish meaningful changes in firing patterns when comparing across two scenarios (*9*), and a no effect label signifies that binned firing rates of a target neuron remained well within the distribution of those measured pre-stimulation, at least for most bins. Under DD, inputs with larger conductances and decay time constants again promoted inhibitory responses. To match experimental results under DD conditions (*10*), in our simulated DD scenario we decreased the decay constants of striatal inputs, and we found that more SNr cells exhibited excitatory or no effect responses when striatum was stimulated, with fewer cases of complete inhibition, consistent with experimental findings (*10*).

We also performed a more in-depth analysis, using the WD to compare the distributions of values of 5 parameters computed separately across model SNr neurons showing either no effect, excitatory responses, or any form of inhibitory responses (Figure 9). That analysis showed that in most cases, the strongest distinction between the excitatory and no response distributions was associated with the term we called *MF*_*input*_, which quantifies the extent of the response of the other SNr inputs that synapse on the target neurons. In other words, when GABAergic terminal stimulation induced an excitatory response in an SNr neuron in our model network, the inhibition of the synaptic neighbors of that neuron (i.e., relatively negative *MF*_*input*_, see and Figure 9E and Figure S8E,J) was likely to be an important contributing factor. This same factor emerged as a prediction in another recent computational study (*26*); however, that work included inputs from dSPNs only to a subset of SNr neurons, with most excited SNr neurons not directly inhibited, whereas our work shows that the disinhibition mechanism still applies without this limitation on input. We also found that for striatal stimulation of SNr in the naive case, an elevated strength of reciprocal input from other SNr neurons was associated with excitatory SNr responses. This observation may represent a factor that synergizes with *MF*_*input*_; that is, those SNr neurons that were more strongly inhibited by other SNr cells before stimulation will show a greater relative increase in firing from the inhibition of their neighbors than will those that were only weakly inhibited by those neighbors.

Multiple factors were associated with a higher likelihood of inhibitory responses in SNr neurons. In addition to the conductance associated with the stimulated terminals as discussed above, more hyperpolarized GABA reversal potentials and weaker inhibition from other SNr neurons were also associated with inhibitory responses. The former factor is quite natural, since more negative *E*_*GABA*_ makes GABAergic inputs effectively stronger. Past modeling work suggests that *E*_*GABA*_ could be modulated across the SNr based on ongoing GPe inputs and the associated chloride loading that they induce (*8*). As for the latter, we find that weaker inhibition from other SNr neurons is correlated with higher SNr neuron firing rates prior to terminal stimulation. In theory, such faster baseline firing could provide more opportunity for inhibitory responses to GABAergic inputs. In our prior analysis of these recordings (*10*), we did not find that SNr baseline firing rate predicted response, so the situation is nuanced. Nonetheless, it does appear that SNr neurons that exhibit relatively high firing rates associated with relatively weak collateral SNr input are likely to be inhibited by terminal stimulation.

Almost all of the differences between parameter distributions of SNr neurons exhibiting inhibitory versus no effect responses to striatal stimulation were diminished under DD (Figure 9C), consistent with a moderation of the negative part of the modulation factor distribution (compare Figure 7E vs. Figure 8E). Interestingly, the shorter decay time of striatal inputs under DD is important in limiting their efficacy and in shifting the overall SNr response profile away from inhibitory responses, but this time constant does not show up as an important factor in our WD analysis; thus, although the overall shift in the *τ*_*syn*_ distribution limits the extent of SNr inhibitory responses to striatal stimulation, the particular value of *τ*_*syn*_ within the shifted *τ*_*syn*_ distribution associated with DD does not seem to be an important factor, relative to others, in determining individual SNr neuron responses.

### Assumptions and Limitations

When comparing the simulated models against experimental observations, we only focused on firing rate and CV distributions. Although it would have been reasonable to consider other neuronal or network firing properties, firing rate and CV are fundamental characteristics that can be computed in standard ways from collections of individual unit recordings. Harnessing this information provided us with a challenging yet attainable target for constraining our network simulations and hence proved ideal for this study. A natural topic for future work will be to investigate other properties of activity in the model network that we have constructed.

With our matched neuronal database, we were able to generate 10 slice models out of 300 randomly sampled networks that met our criteria for capturing the data. It is likely that expanding the database of neurons that we used, for example by taking more than just the single best-matching model neuron corresponding to each experimentally recorded neuron, or creating more randomly sampled networks would generate a larger pool of successful slice models. However, the 10 networks that we produced was a helpful sample size to allow us progress to the subsequent, *in vivo* aspects of our study. That is, at each successive stage of our simulations, we performed additional parameter sweeps and extraction of best-performing networks; starting from an overly large sample early on would have resulted in a combinatorial explosion later in the process. Indeed, when performing the *in vivo* parameter sweep for each slice model, we found many successful matches to the *in vivo* data from naive animals and restricted to the three matching models with the lowest relative error for each of the 10, and we implemented a similar restriction process when comparing to DD data later in our study.

We assign each synaptic connection in our network a fixed weight, ignoring possible plasticity effects. We used synaptic conductances and time constants for GPe and striatal inputs onto SNr neurons that were measured in slice preparations (*10*) but tuned down the strengths of the GPe synapses onto our model SNr neurons by multiplication of all conductances by 1/10. Since significant synaptic depression of GPe synapses has been reported (*35, 36*) and GPe neurons have a high baseline firing rate, it is reasonable that their synapses function in a depressed state in *i*n vivo conditions. Since the data to which we compare our model when incorporating GPe stimulation was indeed recorded *i*n vivo, we can consider this scaling as a synaptic depression factor. The model could be strengthened if this factor were measured precisely or if synaptic plasticity at the level of vesicle dynamics were modeled and included, although that step would necessitate significant additional complexity and parameter fitting. An assumption that we made about the striatal synaptic connections in our model was the introduction of an anti-correlation between the strengths of striatal inputs to model SNr neurons and the GABA reversal potentials at the dendrites where these synapses occur. We found that this feature improved our model fit to experimental data on SNr responses to striatal terminal stimulation, and hence this relationship represents a prediction of this modeling work (Figure S7). Finally, regarding the SNr-to-SNr connections in our model, we imposed the sparse connectivity reported in the literature (*16*) but did not include any spatial dependence or patterning in the connection strengths, which may be an oversimplification.

We introduced stochasticity in the background input strengths in our model, hoping to capture the most prominent noise sources within the SNr network. This stochasticity was modeled using OU processes; although this is a standard approach in neuroscience modeling, whether our stochastic processes exhibit the correct parameter tuning and other properties remains uncertain without experimental verification; moreover, for the inputs from STN, this approach necessarily neglects any temporal patterning such as oscillations or bursts that may occur in the DD state (*37,38*). Finally, in our analysis of model network performance, we compared simulated network dynamics against single unit experimental recordings collected across multiple animals. Of course, the recorded units were embedded within biological networks, but we cannot be certain that the distributions of firing properties assembled in this way match those that would be present across a connected network.

Overall, recent results paint a complicated picture of how SNr neurons respond to GABAergic inputs and how their activity and responses change under DD. In general, SNr activity appears to be relatively mildly affected by GPe inputs and more strongly modulated by striatal inputs (*10*) (Figure 7), but with a subset of excitatory responses in both cases that would not be expected based on the inhibitory properties of GABA (*7, 9, 10*). Modeling work has suggested that the diversity of responses may relate to heterogeneous GABA reversal potentials across SNr neurons, perhaps related to their chloride load as tuned by ongoing GPe activity (*8*). Under DD, effective synaptic inputs from striatal neurons to SNr weaken through a shortening of their decay time and are less likely to completely inhibit SNr neurons (*10*). On the other hand, lower rates of GPe firing in DD could result in a hyperpolarizing shift in *E*_*GABA*_ that would make at least some inhibitory inputs more effective (*8*). The results of our present study support the idea that TRPC3 channels on dendrites of SNr neurons weaken under DD, which would lower SNr neuronal excitability. These last two sets of results are consistent with the shift of the SNr firing rate distribution towards lower levels under DD (Figure S2D). As our results on SNr response modulation show, however, because SNr neurons comprise a network that is synaptically interconnected, albeit sparsely, reductions in activity in some SNr neurons can result in increased firing of others. Therefore, when combined, these factors set the stage for more variability in SNr firing under DD, consistent with the upward shift in the CV distribution in our data (Figure S2E), with neurons potentially taking turns being suppressed or active. Moreover, these conditions of an altered balance of inhibitory inputs together with weakened intrinsic neuronal dynamics would be conducive to the emergence of oscillations within the SNr network, as are associated with the DD state and parkinsonism.

## Materials and Methods

### Single-Neuron SNr Model

The model used in this work is an extension of the two-compartment (soma and dendrite) SNr model from (*8*). The earlier model included GABAergic inputs from GPe and Str stimulation as well as a static GABA conductance, representing ongoing background or tonic inputs from GPe and Str, that was taken into account specifically in modeling chloride dynamics in each compartment. We have treated these background components more realistically by including the appropriate term in the voltage dynamics of each compartment and allowing each conductance, denoted by 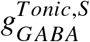 and 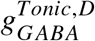, to vary dynamically as a stochastic process. The inclusion of these background synaptic dynamics required us to determine new ranges for the KCC2 pump conductances and mean background GABA conductances such that the range of GABA reversal potentials were still similar to the previous version of the model (*8*). The governing equations for the somatic (*V*_*S*_) and dendritic (*V*_*D*_) voltage dynamics (Eq. 1 and Eq. 2 in (*8*), respectively) have been updated to include the synaptic conductance fluctuations as well as stochastic, ongoing STN inputs to the dendritic compartment and time-varying channel noise to both compartments (lumped into a single term rather than subdivided into terms for specific channel types) and now take the form

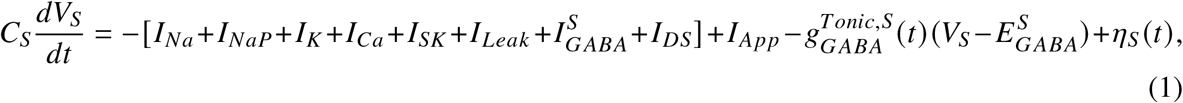

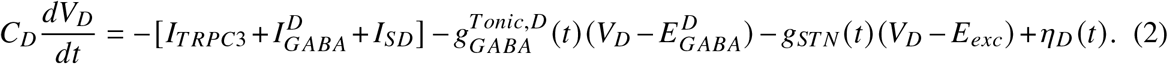

The stochastically-varying conductances are modeled as Ornstein-Uhlenbeck (OU) (*17*) processes:

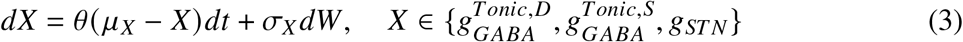

where each *μ*_*X*_ is a mean to which a deterministic component of the conductance dynamics reverts at a rate *θ* and each *σ*_*X*_ is the standard deviation of a stochastic Wiener or white noise process. Similarly, channel fluctuations are denoted by *η*_*S*_ (*t*) and *η*_*D*_ (*t*) and obey

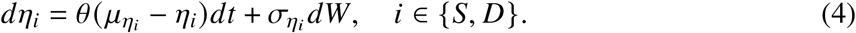

Note that all stochastic terms in the model are independent. Values of newly introduced parameters and those changed relative to the earlier, published version of the model are listed in Table 1. With these values, the baseline model firing rate is approximately 10 Hz when *I*_*App*_ = 0.

### SNr Network Model

Following the earlier work (*8*), we consider coupled networks of 100 SNr neurons. We tuned parameters for these networks to replicate various experimental recordings, starting with data collected in slices and then progressing to *in vivo* data from a variety of conditions (Figure 1).

To construct model SNr networks based on experimental observations from slice preparations, we synaptically coupled 100 model SNr neurons; since the literature reports an average of 1.16 connections per each of 81 SNr units (*16*), we coupled them with a pairwise connection probability of approximately 0.014. Synaptic conductances were randomly and independently selected from a normal distribution, 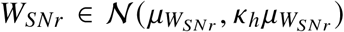. We included background GABAergic inputs with strength 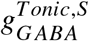 in the dynamics of model neuron somatic compartment voltages (Eq. (1)). As with other conductances in the model, heterogeneity was introduced into the network by selecting values for individual model neurons from a normal distribution, N, around a fixed mean (given from (*8*) for the parameters also used there), such that

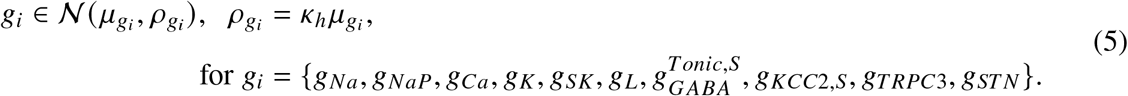

For the dendrites, we introduced a scaling factor to reflect the appropriate compartment capacitance values as given in the following equation:

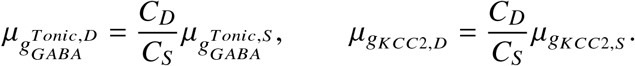

We set the rate of reversion to the mean in equation (5), as θ = 0.001. No STN dynamics were included in the model in the slice condition. We did, however, include channel noise in each model neuron compartment, as described in equation (4), with θ = 0.1 and 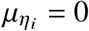 in both cases. Finally, we kept most model reversal potentials fixed, except that for each model neuron we randomly selected 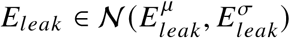 with 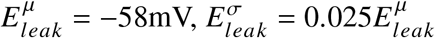.

To try to tune the model to fit the data, we varied the following parameters:

1. heterogeneity factor, *k*_*h*_ ∈ {0.1, 0.2, 0.3, 0.4}, from equation (5)
2. SNr-to-SNr mean synaptic strength, 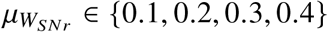
3. mean somatic KCC2 pump conductance,

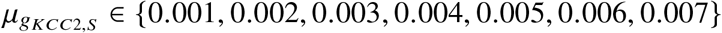
4. mean strength of background GABAergic inputs to the soma,

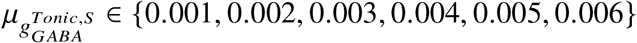
5. channel noise volatility, 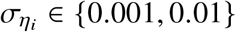

These options together comprise 1,344 parameter combinations that we considered for the network model.

#### Fitting to slice data

Out of the 1,344 slice model simulations, no single model passed all of the criteria for matching to the experimental data that we applied. Within models, specific neurons exhibited firing properties similar to experimental observations of individual units, however. Thus, we constructed a database with all 134,400 neurons from the 1,344 slice models and compared the firing properties of each model neuron with the experimental recordings. Specifically, we iterated through all *n* = 252 experimental observations from slice preparations and, for each, we identified the model unit from the 134,400 neuron database that was the closest match, determined as the model unit with the minimum of the average of the relative errors between its firing rate and CV and those of the experimental neuron.

This selection process yielded a set of model neurons, χ_*slice*_, matching the experimental data. From this set, we randomly selected the 100 units to use in our model networks. Specifically, we sliced the firing rate histogram from the recorded neurons into *B* bins of 5 Hz each, {*b*_*i*_ : *i* = 1,.. ., *B*}, computed the proportion of experimental units in each bin (*p*_*i*_ = #(*b*_*i*_)/252), and, for each *i*, randomly selected *p*_*i*_ units from those in χ_*slice*_ with firing rates in the appropriate 5 Hz range, to obtain a model network with firing rate proportions that matched the data. Note that in cases when we did not have at least *p*_*i*_ units in χ_*slice*_ for some *i*, we randomly selected extra units from all of χ_*slice*_ to ensure that we had 100 total model neurons in our model network.

We repeated this process for each of 300 random seeds to obtain 300 distinct model neuron networks in a way that maintained agreement in the distribution of firing rates between the neurons comprising the model network and the experimentally recorded neurons. For each selected collection of 100 model neurons, we introduced reciprocal SNr synapses with a mean *W*_*SNr*_ value defined as the average of the 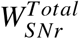 divided by the average of the number of connections from the sampled neurons and a probability defined as the average number of connections divided by 100 times the size of the network, computed from the sampled neurons. Out of the 300 tuned model networks, 10 were not statistically significant different from the data in terms of the overall firing rate distribution (K-S test, *p* > 0.05), mean firing rate (*t*-test, *p* > 0.05), and mean CV (*t*-test, *p* > 0.05). All networks were statistically significantly different when comparing full CV distributions with experimental observations from slice preparations.

#### *Parameter updates for* in vivo *networks*

We built upon the 10 model networks tuned to slice conditions to create *in vivo* model networks. In the *in vivo* setting, since we were modeling an intact SNr, we doubled the SNr-SNr synaptic connection probability from the slice condition to 0.03, increased the current noise volatility 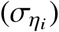, increased the mean strength and volatility of the background/tonic GABAergic inputs, and included an excitatory tonic input channel to represent inputs to SNr from the STN. We swept through scaling factors *k*_*i*_ where *i* are the parameters associated with these *in vivo* features to produce and identify model networks that match the *in vivo* experimental observations:

1. 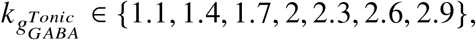
2. 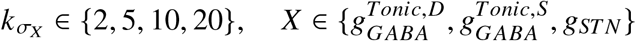
3. 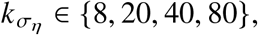
4. 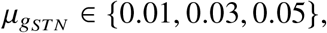
5. 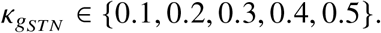

Recall that the STN synaptic conductance to each neuron follows an OU process (Eq. 3) with *θ* = 0.001 and the noise volatility of the STN OU process as 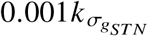. Collectively, these parameter options result in 1680 combinations for the *in vivo* setting, yielding 16,800 simulations when applied to each of the 10 slice models. Out of the 16,800 simulations, we found 930 *in vivo* models that were not statistically different when compared with *in vivo* experimental observations for mean firing rate (K-S test, *p* > 0.05), firing rate distribution (*t*-test, *p* > 0.05), mean CV (K-S test, *p* > 0.05), and CV distribution (*t*-test, *p* > 0.05).

For each of these 930 models, we compared the distributions for the simulated and recorded firing rate and CV by defining the relative error in the firing rate (Rel. Error FR) and and relative error in the CV (Rel. Error CV) as the following, where *i* and *k* represent the bin heights in the model histogram and the experimental histogram, respectively:

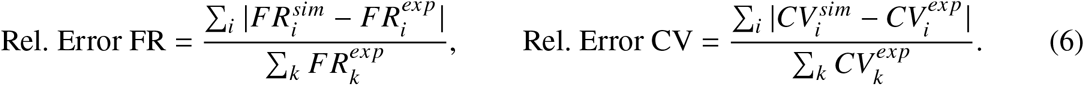

For our analysis under *in vivo* conditions, we kept the three models with the lowest average errors, defined by averaging the quantities from Eq. 6, generated from each slice model. These models were also used to simulate 20Hz pulse stimulation (implementation described in the next section) and to model changes under dopamine depletion (DD).

### Stimulation of GABAergic Terminals in SNr

Similarly to (*8*), we simulated stimulation of GABAergic terminals impinging on the SNr from the direct (D1-SPN neurons in Str) and from the indirect (GPe) pathways. More specifically, to represent pulsatile stimulation of these terminals, we introduced a rise in synaptic current that only lasted for one time step which were then subject to synaptic decay as described in (*8*). Stimulation was simulated as 10 trials of 20 Hz pulses, as used experimentally (*10*); these were run for one second each. Altogether, simulations of stimulation experiments comprised 35 simulated seconds: the first 5 seconds corresponded to baseline SNr activity and the subsequent 30 seconds spanned 10 stimulation trials, with one second each in the pre-stimulation, stimulation, and post-stimulation time periods.

The equations governing the indirect and direct pathway inputs to SNr were the same as in past work (*8*). However, the values for the synaptic decay constants and direct/indirect-to-SNr connection strengths were based on our recent experimental work (*10*). We used that data to generate a linear regression of decay constant versus direct/indirect-to-SNr connection strengths. Then, we selected the connection strengths uniformly at random from the experimentally derived range and for each, we randomly selected the decay constant from a 99% confidence interval around the linear regression line at that connection strength value. For the indirect pathway, we multiplied all GPe input weights, denoted by *W*_*GPe*_, by a scaling factor of 1/10 (perhaps representing depression of GPe synapses present *in vivo*; see *Discussion*), which improved model agreement with experimental results, whereas no scaling factor was applied to striatal input weights, *W*_*Str*_ . We repeated the stimulation simulations three times – corresponding to different instantiations of the random values for decay constants and direct/indirect-to-SNr connection strengths – for each of the top three *in vivo* models derived from each slice model (30 models total), which resulted in 90 model network simulations of stimulation experiments.

### Dopamine Depleted *in vivo* SNr Model

Experimental recordings showed significant differences in both baseline activity and responses to stimulation for SNr neurons when comparing naive and dopamine depleted conditions (*10*). In our model, eight parameters were considered to potentially change under dopamine depletion, based on previous experimental results (*4, 5, 10, 22–25*). The parameters theorized to possibly decrease were the conductances 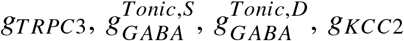 for the somatic and dendritic compartments, and *g*_*TRPC*3_. The parameters considered to possibly increase were the current noise volatility, 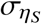 and 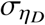, and the STN input strength *g*_*STN*_ . This gives 7 total parameters that we varied from a baseline value of 1, using independent scaling factors (the count is 7 and not 8 because current noise volatility was scaled by the same factor in the somatic and dendritic compartments).

As for the scaling factors, each parameter *i* for which we explored reductions in value was scaled by each of the factors *k*_*i*_ ∈ {0.33, 0.66, 1}, meaning a strongly reduced scaling (0.33), a moderately reduced scaling (0.66), and a baseline scaling (1). The current noise volatility was scaled by 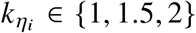 for *i* ∈ *S, D* and the the STN conductance was scaled by 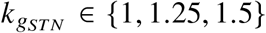. With 7 parameters each being set to each of three values, we considered 2,187 possible parameter change combinations for each network. We specifically identified the simulations that matched DD data based on the same matching criteria as we used in the baseline or naive *in vivo* case (*p*-value for the Kolmogrov-Smirnov test greater than 0.05 and *p*-value for a *t*-test greater than 0.05 for comparisons between experimental and simulated firing rates and CV distributions). For each *in vivo* model, we kept the 3 DD versions with the lowest average errors relative to the stimulation experiments in DD conditions (Eq. 6).

### Simulation and Data Analysis

Simulated and experimentally recorded SNr responses to stimulation of input pathways were classified using the STReaC toolbox (*9*). The SNr model was built in C++. Parameter selections and data analysis were performed in Python (version 3.12.11). All simulations and analysis were performed on a 2020 13-inch MacBook Pro with an Apple M1 chip. See Data Availability statement for details on the availability of relevant code needed to reproduce results found in this work.

### Experimental Methods

For collection of slice data, we used acute brain slices from transgenic mice to perform cell-attached recordings of spontaneous firing in SNr neurons. The *in vivo* data used for this study, including relevant experimental methods, are described in detail in a previous study (*10*). In brief, *in vivo* data was collected from adult male and female mice greater than 8 weeks old. Data from dopamine depleted mice were collected 3-5 days after bilateral 6-OHDA injections in the medial forebrain bundle (MFB), as described in (*39*). The optogenetic protocol for D1-MSN and GPe-PV axon terminal stimulation in the SNr was 5ms light pulses at 20Hz for 10 seconds, 1 mW light intensity (blue light, 465 nm wavelength) and we focused only on the first second of stimulation. In some cases, we used multiple baseline recordings from the same unit that were taken at different times. We verified that the differences between firing rates and CVs of recordings from the same unit were not different from those for randomly selected units.

### Artificial Intelligence Tools

We used ChatGPT-4o to adjust Python code for the decision trees in Figure 6 and Figure S4. Example prompts are “Adjust the code for this decision tree such that a directional color-coded tree is generated based on conditional good fit rates, make the branches have ‘True’ or ‘False’ labeling, with square nodes” and “Adjust the code for this figure generation such that it is a horizontal layout showing all paths with arrows for edges.” Initial code and original decision trees were refined with ChatGPT-4o’s output to find finalized layouts and visual design. Other instances of usage include debugging with a common prompt of “Find why this error is occurring.” Generative responses were then examined for errors and adjusted appropriately.

## Acknowledgments

We thank Ryan S. Phillips (Children’s Hospital, Seattle) for initial groundwork with the SNr model. ChatGPT-4o was used to assist with code regarding certain figure layouts and with debugging.

## Author contributions

Conceptualization: JEP, JER. Methodology: JEP, JER. Investigation: JEP, JER. Formal analysis: JEP, JER. Visualization: JEP, JER. Data curation: JEP, AA, YAG. Validation: JEP, JER. Software: JEP. Project administration: AHG, JER. Funding acquisition: AHG, JER. Resources: JEP, AA, YEG, AHG, JER. Supervision: AHG, JER. Writing—original draft: JEP, JER. Writing—review and editing: JEP, AA, YEG, AHG, JER.

## Data, Code, and Materials Availability

All data and code to replicate is hosted at the following GitHub repository https://github.com/jparker25/Parker_et_al_SNr_network which will be made publicly available upon acceptance.

