## Supplemental figures. for "Altered SNr activity under dopamine depletion: mechanistic predictions from data-driven network models"

**This PDF file includes:**

Figures S1 to S8

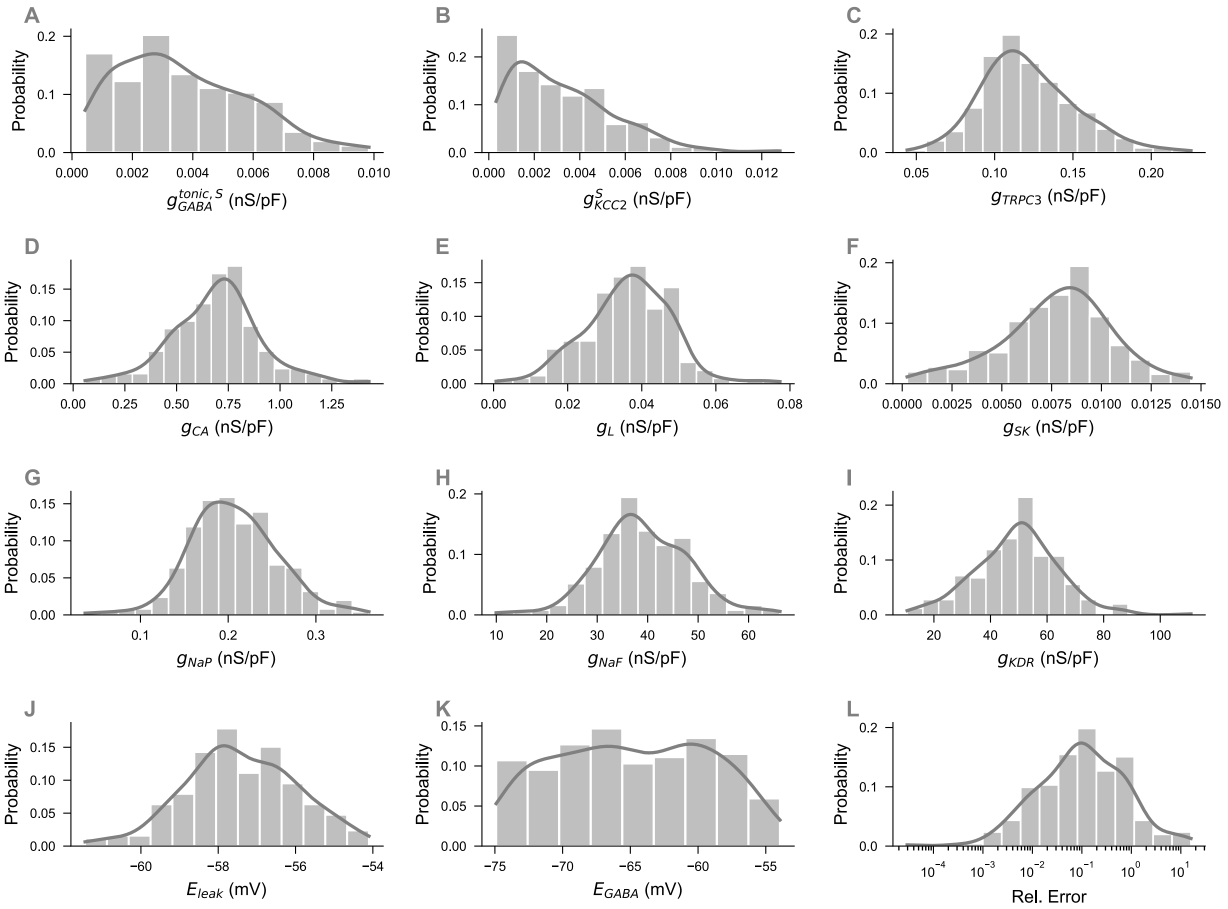


**Figure S1**: **Parameter distributions from individual neurons from across all models that match slice data** (*n* = 10)**.** For each distribution, we show a histogram of the probabilities with which the binned sets of values occur as well a curve representing a kernel density estimate (KDE) of the probability density.

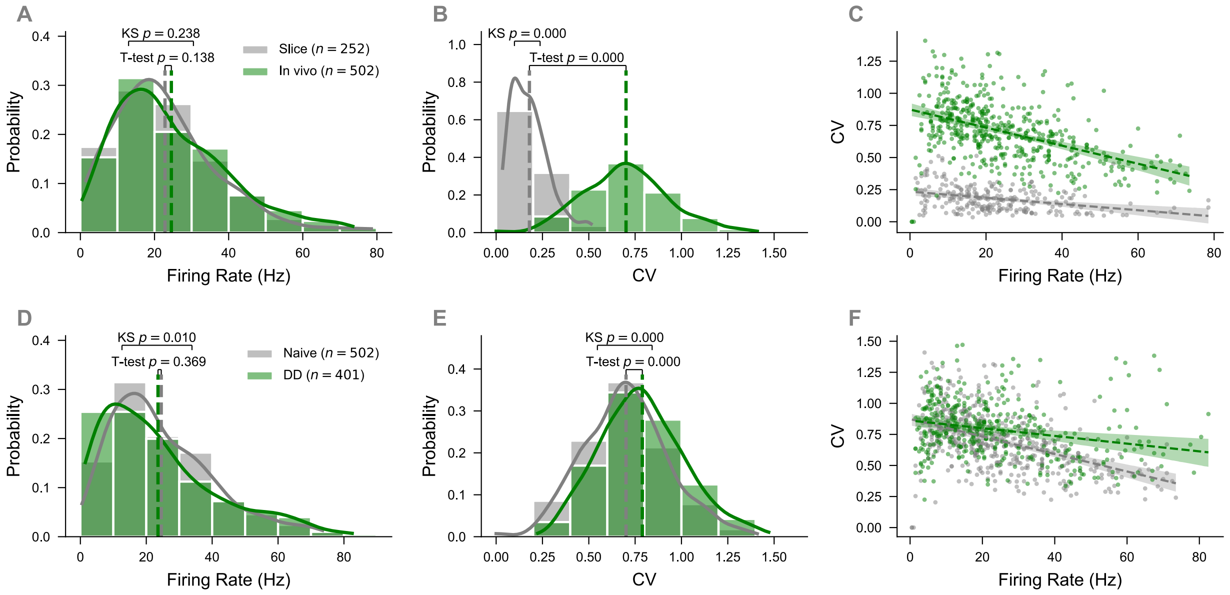


**Figure S2**: **Comparison of experimentally measured firing rate and CV in naive slice and *in vivo* and dopamine depleted *in vivo* settings.** (**A**) Naive firing rate histogram in slice (gray) versus *in vivo* (green). Distributions of firing rates were not significantly different (K-S test, $p = 0.238$) nor were the mean rates ($t$-test, $p = 0.138$). (**B**) Naive CV histogram in slice (gray) versus *in vivo* (green). Distribution (K-S test, $p < 0.001$) and means ($t$-test, $p < 0.001$) of CVs differed significantly. (**C**) Linear regression with 99% confidence intervals comparing *in vivo* (green) and slice (gray) data. The regression line for the *in vivo* data has slope = $-0.007$, intercept = 0.874, and $R^2 = 0.220$ while the regression line for the slice data has slope = $-0.002$, intercept = 0.235, and $R^2 = 0.118$. (**D**) Firing rate histogram for naive *in vivo* (gray) versus dopamine depleted *in vivo* (green) data. Distributions of firing rates were significantly different (K-S test, $p < 0.05$) but the mean rates were not ($t$-test, $p = 0.369$). (**E**) CV histogram for naive *in vivo* (gray) versus dopamine depleted *in vivo* (green) data. Distributions (K-S test, $p < 0.001$) and means ($t$-test, $p < 0.001$) of CVs differed significantly. (**F**) Linear regression with 99% confidence intervals comparing naive *in vivo* (gray) and dopamine depleted *in vivo* (green) data. Naive *in vivo* regression line has slope = $-0.007$, intercept = 0.874, and $R^2 = 0.220$ while DD *in vivo* regression line has slope = $-0.003$, intercept = 0.862, and $R^2 = 0.059$.

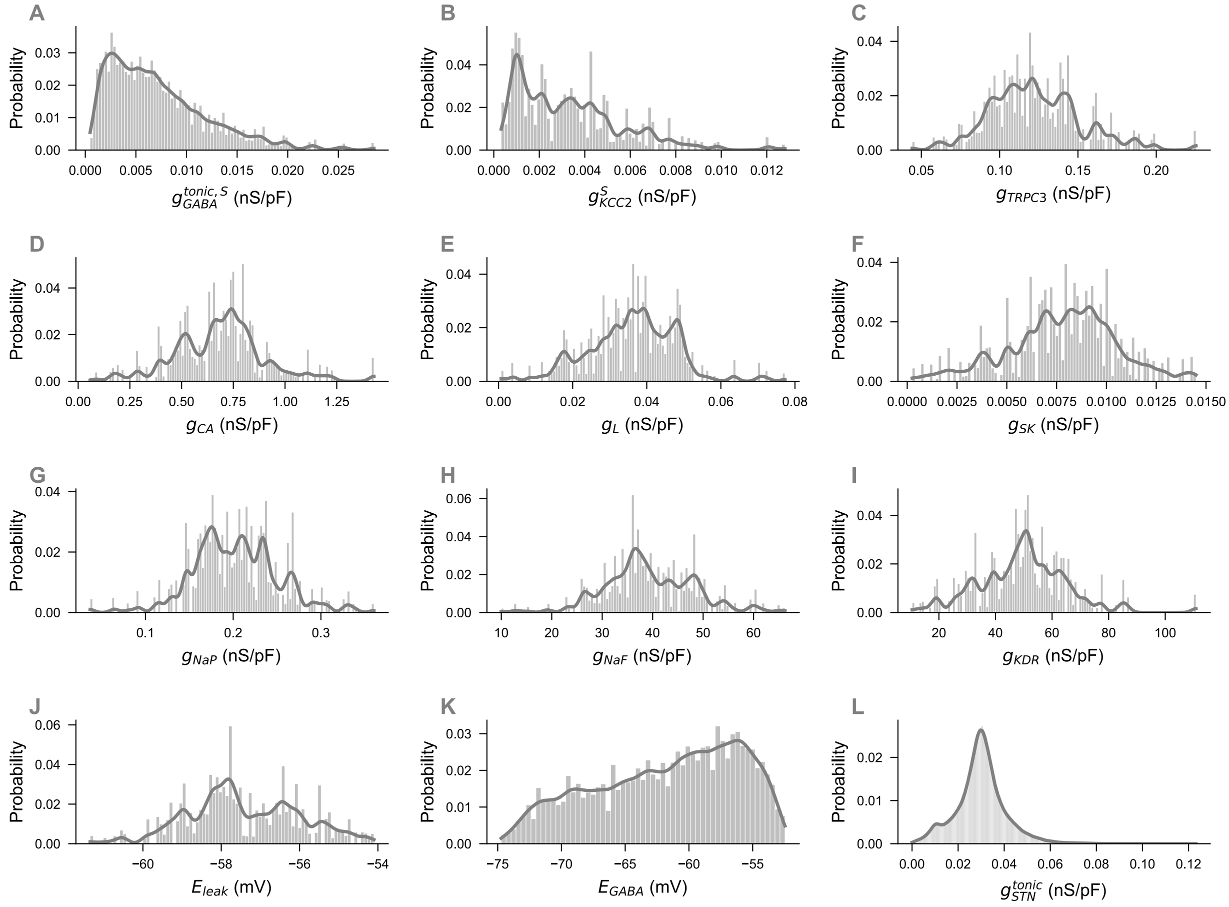


**Figure S3**: **Parameter distributions from individual neurons from across all models that match *in vivo* data** ($n$ = 930). For each distribution, we show a histogram of the probabilities with which the binned sets of values occur as well a curve representing a kernel density estimate (KDE) of the probability density.

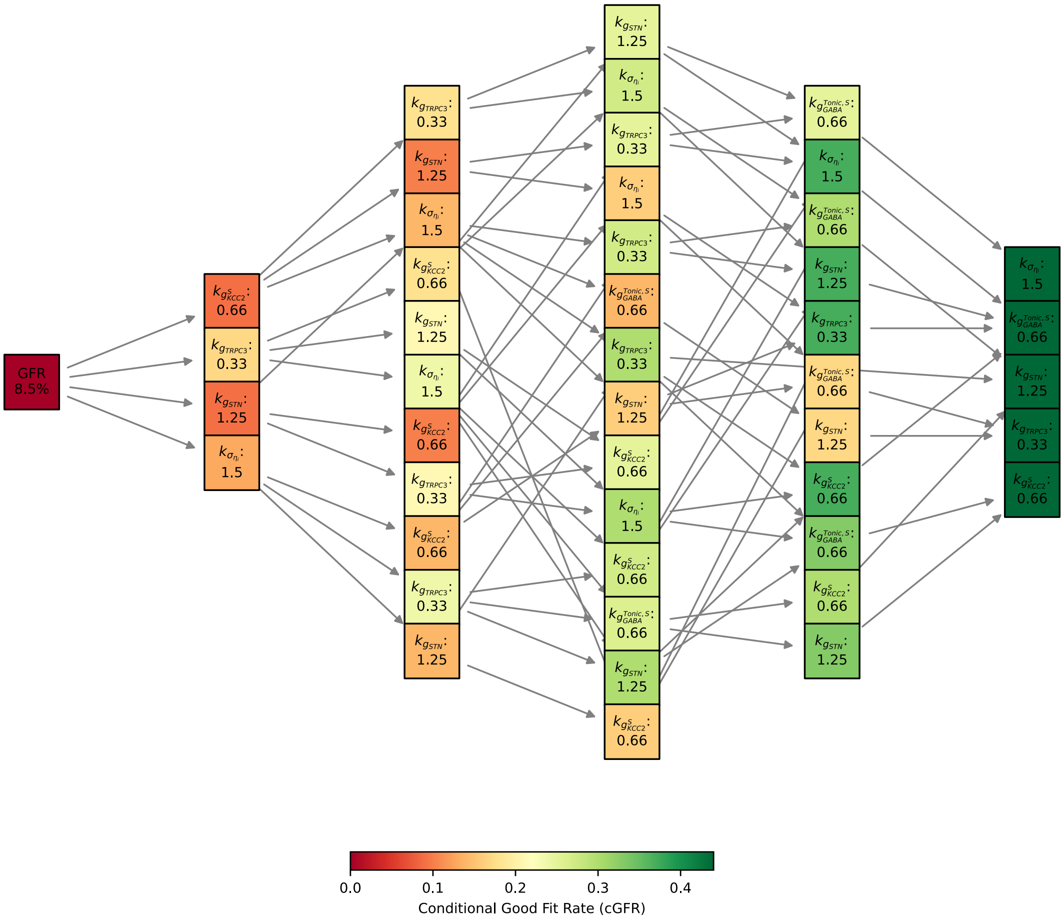


**Figure S4**: **Decision tree for dopamine depleted changes.** A decision tree improving the conditional good fit rate (rounded to two decimals) at each level. Only paths where a change (e.g. $k_i \neq 1.0$) were allowed.

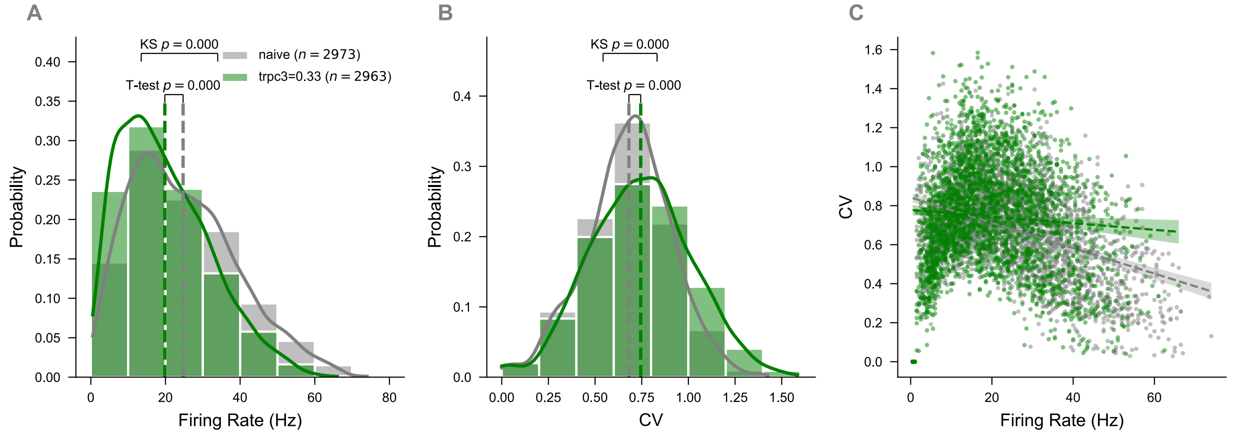


**Figure S5**: **Firing rate statistics for naive *in vivo* models versus models with $k_{TRPC3} = 0.33$.** (**A**) Firing rate histogram in naive models (gray) versus $k_{TRPC3} = 0.33$ models (green). Distribution (K-S test, $p < 0.001$)and mean values ($t$-test, $p < 0.001$) of firing rates differed significantly. (**B**) CV histogram in naive models (gray) versus $k_{TRPC3} = 0.33$ models (green). Distribution (K-S test, $p < 0.001$) and means ($t$-test, $p < 0.001$) of CVs differed significantly. (**C**) Linear regression with 99% confident intervals comparing naive models (gray) and $k_{TRPC3} = 0.33$ models (green). Naive models linear regression has slope = $-0.007$, intercept = 0.842, and $R^2 = 0.158$ while $k_{TRPC3} = 0.33$ models linear regression has slope = $-0.002$, intercept = 0.778, and $R^2 = 0.006$.

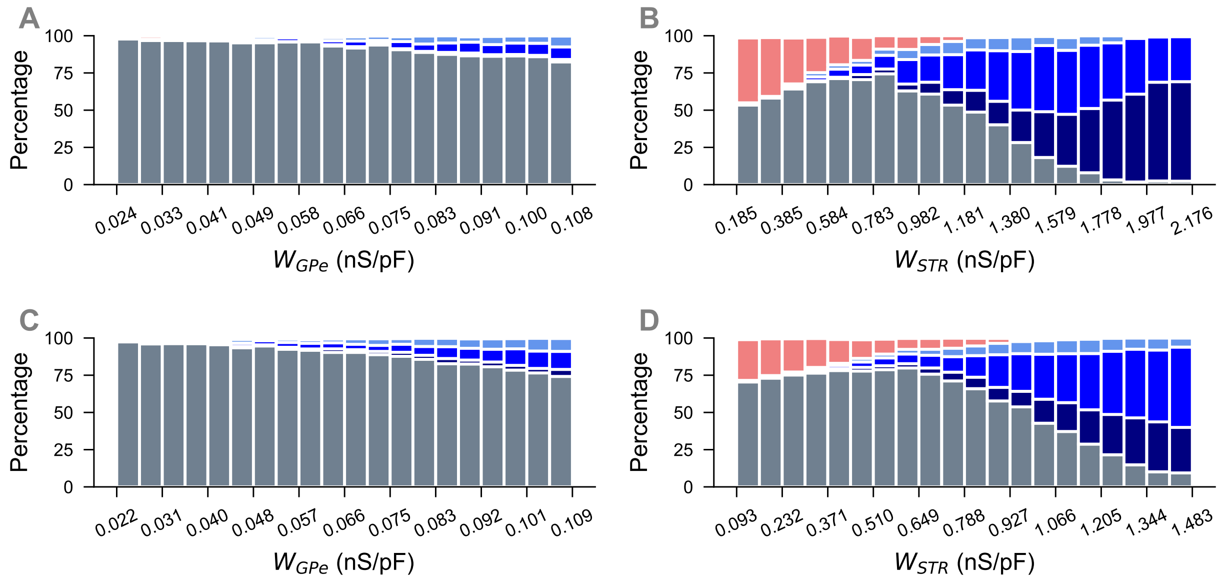


**Figure S6**: **Dependence of SNr response distribution on input weights.** (**A,B**) Responses to (A) GPe and (B) Str stimulation in control conditions from Figure 7D and Figure 7H, respectively, separated by equal bins of $W_{GPe}$ and $W_{Str}$. (**C,D**) Responses to (C) GPe and (D) Str stimulation in DD conditions from Figure 8D and Figure 8H, respectively, separated by equal bins of $W_{GPe}$ and $W_{Str}$. Colors correspond to the response types identified by the STReaC toolbox (*9*).

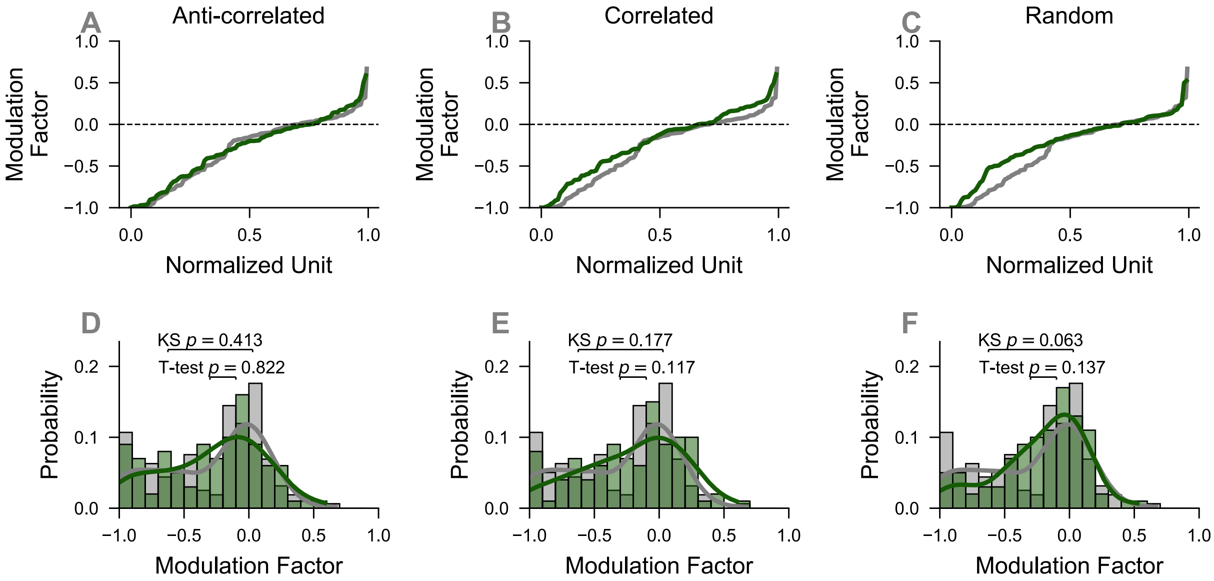


**Figure S7**: **Model responses to stimulation depend on $W_{Str}$ relationship with $E^D_{GABA}$.** (**A**-**C**) Normalized cell number versus modulation factor (MF) for experimental responses (grey) and model responses (green) to naive D1 stimulation with anti-correlated (panel (**A**)), correlated (panel (**B**)), and random (panel (**C**)) relationships between $W_{Str}$ and $E^D_{GABA}$. (**D**-**F**) Histograms of modulation factors corresponding to panels (**A**) - (**C**), respectively. MF distributions and means statistically agree with experimental data for anti-correlated (panel (**D**), $MF_{model} = -0.282 \pm 0.387$, $MF_{exp} = -0.293 \pm 0.396$, K-S test $p = 0.413$, T-test $p = 0.822$), correlated (panel (**E**), $MF_{model} = -0.214 \pm 0.384$, $MF_{exp} = -0.293 \pm 0.396$, K-S test $p = 0.177$, T-test $p = 0.117$), and random (panel (**F**), $MF_{model} = -0.222 \pm 0.336$, $MF_{exp} = -0.293 \pm 0.396$, K-S test $p = 0.063$, T-test $p = 0.137$) relationships, with strongest agreement for the anti-correlated case.

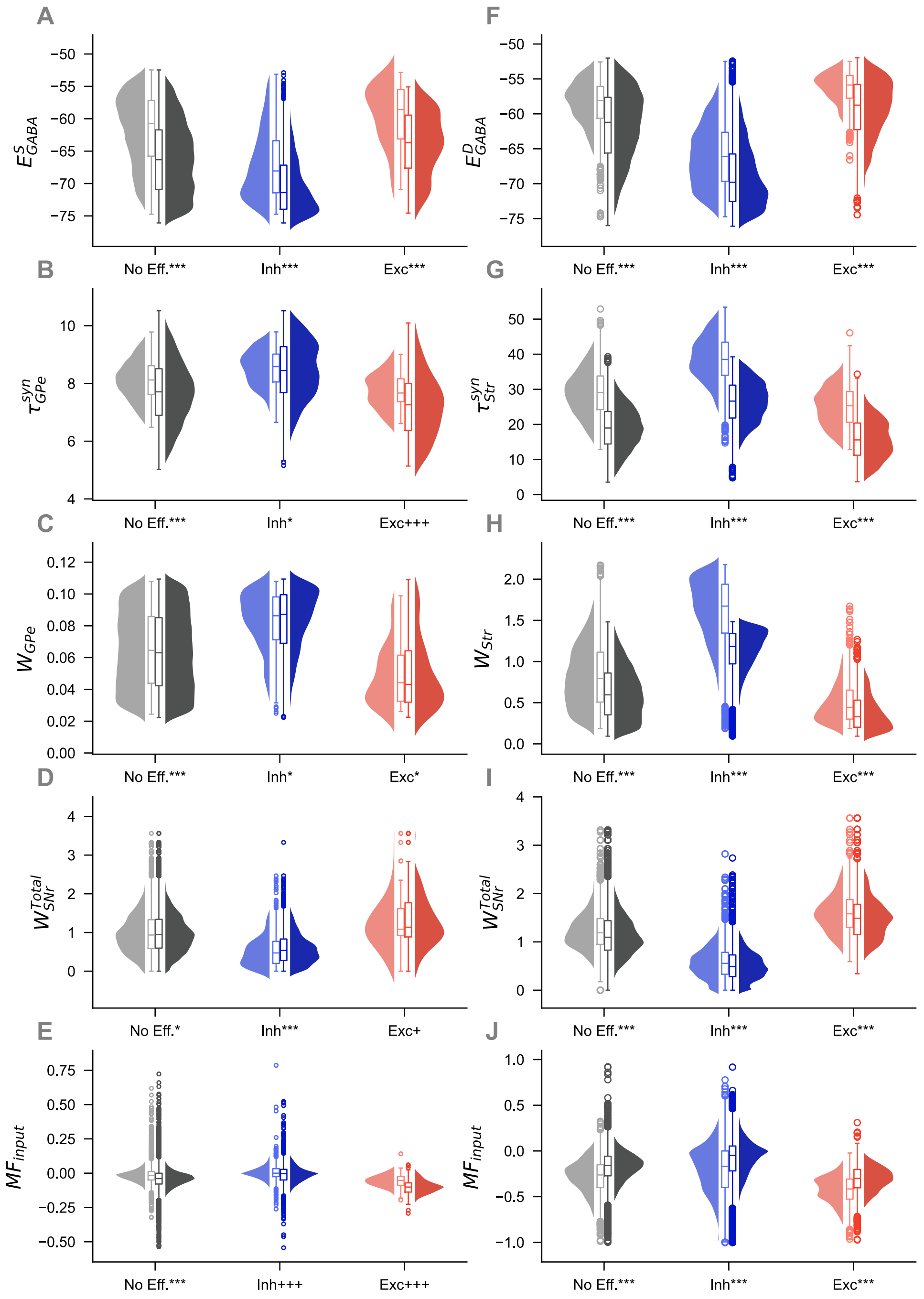


**Figure S8**: **Violin plots illustrating the distributions of measured parameters in the GPe (left column) and Str (right column).(A,B)** $E_{GABA}$ values. (**C,D**) Synaptic decay constants, $\tau^{syn}_{GPe}$. (**E,F**) Synaptic weights for stimulation. (**G,H**) Total network synaptic input. (**I,J**) Average modulation factor for input cells. Asterisks (crosses) indicate statistical differences between naive (left) and diseased (right) via Mann Whitney U test ($t$-test). Significance levels are $^{***}p < 0.001$, $^{**}p < 0.01$, $^{*}p < 0.05$ (analogous for crosses).
